# Systemic influences of mammary cancer on monocytes in mice

**DOI:** 10.1101/2021.12.24.474104

**Authors:** A Robinson, M Burgess, S Webb, PA Louwe, Z Ouyang, D Skola, CZ Han, NN Batada, V González-Huici, L Cassetta, CK Glass, SJ Jenkins, JW Pollard

**Affiliations:** Centre for Reproductive Health, Queen’s Medical Research Institute, University of Edinburgh, UK; Centre for Inflammation Research, Queen’s Medical Research Institute, University of Edinburgh, UK; Department of Cellular & Molecular Medicine, University of San Diego, USA; Institute of Genetic and Molecular Medicine, University of Edinburgh, UK

**Keywords:** Breast cancer, monocytes, mouse, human, myeloid

## Abstract

There is a growing body of evidence that cancer causes systemic changes. These influences are most evident in the bone marrow and blood, particularly the myeloid compartment. Here we show using mouse models of breast cancer caused by the mammary epithelial expression of the Polyoma middle T antigen that there is an increase in the number of circulating and splenic monocytes. In the circulation, cancer does not affect ratios of classical to non-classical populations monocytes nor their halflives. Single cell RNA sequencing also indicates that cancer does not induce any new monocyte populations. In the bone marrow cancer does not change monocytic progenitor number is unaffected but the proliferation rate of monocytes is higher thus providing an explanation for expansion in the circulating number. Deep RNA sequencing of these monocytic populations reveals cancer causes changes in the classical monocyte compartment with changes evident in bone marrow monocytes but more in the blood suggesting influences in both compartments. Down regulation of interferon type 1 signalling and antigen presentation were the most prominent. Consistent with this analysis down regulated genes are enriched with STAT1/STAT2 binding sites in their promoter, transcription factors required for type 1 interferon signalling. However, these transcriptome changes in mice did not replicate those found in patients with breast cancer. Consequently, mouse models of cancer may be insufficient to study the systemic influences of human cancer.

## Introduction

Monocytes are key players in the innate immune system, surveying the vasculature in the steady state or being recruited to normal tissues and to sites of infection or tissue damage where they terminally differentiate into macrophages and dendritic cells [1]. The current consensus is that blood monocytes largely derive in the bone marrow (BM) from hematopoietic stems cells (HSC). These progenitors differentiate through several steps to give the restricted erythro-myeloid progenitors that are present in the more abundant Lin^-^Sca-1 ^-^c-Kit^hi^ (LK) population that in turn can be further sub-divided into the restricted Granulocyte/monocyte (GMP) progenitor. This bi-potent progenitor then differentiates to a macrophage dendritic cell progenitor (MDP) and subsequently, the unipotent monocyte progenitor (cMoP) [2]. However, recent evidence suggests there may be alternative pathways directly from the GMP or even directly from HSCs [3, 4]. Newly formed pre-monocytes proliferate within the BM [5] before being released into the peripheral circulation in a CCR2 dependent fashion [6]. In mice and humans there are two dominant blood populations, the classical (Ly6C^hi^ CCR2^hi^ and CD14^hi^ CCR2^hi^ CD16^-^ respectively) and non-classical populations (Ly6C^lo^ CCR2^-^ and CD16^hi^ CCR2^-^ CD14^dim^ respectively) [7, 8] with an intermediate population with mixed markers (Ly6C^mid^ and CD14^hi^CD16^hi^ respectively) [9, 10]. Studies determining half-lives through pulse labelling DNA synthesis with nucleotide analogues indicate the classical population gives rise to the non-classical population through this intermediate population in both mouse and human [11, 12]. However, the ratio of monocytes in steady state conditions is different between mouse and humans, with the former having almost equal numbers of the two populations whereas the classical subset predominates in humans.

The frequency, activity and fate of monocyte populations has been linked to many diseases, including autoimmunity, chronic inflammation, cardiovascular diseases, and cancer [13–18]. In cancer, classical monocytes are in many cases the source of tumour associated macrophages [19–21] and metastasis associated macrophages [22, 23] that can promote primary tumor progression and metastases through a number of different mechanisms [14, 24–26]. However, in some cases, such as in models of pancreatic and glioblastoma there can also be significant recruitment from yolk sac derived tissue resident macrophages [27, 28]. The specific tumour type within a tissue, for example as shown comparing metastatic brain tumours versus primary brain tumours [28], reinforces the idea that intrinsic tumour features drive the tumour microenvironment landscape [29]. In mouse models of breast cancer, the origin of tumour and metastasis associated macrophages is from circulating monocytes [19, 21] that are recruited via CCL2 synthesised at least in part by tumor cells [19, 23].

There is also growing evidence that cancer has systemic influences on monocytes. For example, primary tumors exert systemic influences to generate the so-called premetastatic niche that enhances metastatic cell seeding at these distant sites [30–32]. The formation of this niche appears in part, through effects in the BM and the subsequent mobilisation of cells and their recruitment to tissue specific sites. These cells have been referred to as “immature” myeloid cells that are probably monocytes and their immediate post-differentiation stages that enhance metastatic cell trophism and seeding [32–35]. Several factors have been shown to be responsible for the formation of pre-metastatic niches including fragments of extracellular matrix (ECM), various ECM modifying enzymes such as LOX1, growth factors like VEGFA and tumor derived exosomes [30] [32].

In several different human cancers, the ratio of classical to non-classical monocytes is also altered. For example, recent studies have shown dramatic increases in the non-classical to classical monocyte ratio in breast and endometrial cancer [36], and in breast cancer this increase of non-classical monocytes is negatively associated with tumor size and pathological stage [37]. In addition, monocytes isolated from patients with cancer have been shown to be transcriptionally altered [36, 38, 39]. These transcriptional alterations are profound enough to drive gene expression signatures that predict the presence of cancer [13, 39]. However, whether similar changes occur in mouse models is unknown.

The PyMT transgenic mouse model (PyMT), in which the polyomavirus middle T protein is expressed under control of the mouse mammary tumour virus promoter whose expression is restricted to the mammary gland epithelium, is a well characterised model for studying ductal breast cancer. In this model, mutifocal tumours develop spontaneously in the mammary glands and evolve in a manner that reflects stages of human ductal cancer [40, 41]. Hence this model has been used to study immunological perturbations that accompany mammary cancer progression. In this model tumour progression leads to the dramatic expansion of circulating Gr1 ^+^ myeloid cells [42]. Focusing predominantly on neutrophils, Casbon et al demonstrated an increase in the production of neutrophils within the BM and an expansion of monocytes within the spleen [42]. However, the use of the traditional anti-Gr1 antibody that detects both Ly6C and Ly6G antigens and thus both neutrophils (Ly6G^+^) and classical monocytes, means that specific cell populations were not assessed in this study. Nevertheless, these findings combined with the findings in humans that circulating monocytes have altered population dynamics and distinct transcriptional signatures in cancer when compared to healthy controls, suggests important functions for monocytes that may be best interrogated in mouse models given the availability of genetic analysis in this species. Hence, we aimed to investigate the effect of tumour progression on the production, turnover, transcriptional signature, and composition of the monocyte compartment in the PyMT model.

## Materials and Methods

### Mice

Male B6Tg(MMTV-PyMT)^634Mul^/^Lellj^ a kind gift from Dr. Sandra J. Gendler (Mayo Clinic, AZ) were bred with female WT or PyMT^-/-^ C57BL/6JCrl mice obtained from within the in house existing colony. 8-week-old C57BL/6JCrl mice as controls were purchased from Charles River Laboratories. FVBg(MMTV-PyMT)^634Mul^ were originally obtained from Dr William Muller and maintained in a colony in-house. Animals were housed and bred under standard conditions of care. All procedures involving mice were conducted in accordance with Arrive guidelines with licensed permission under the UK Animal Scientific Procedures Act (1986) and associated guidelines.

### Flow cytometry and cell sorting

Blood was collected by intra-venous (IV) bleed or cardiac puncture into a syringe containing 0.5M EDTA (Fisher Scientific). Single-cell suspensions were prepared using red blood cell lysis buffer (Biolegend) for blood and red blood cell lysis buffer for BM (MERCK) and then filtering with a 70μ filter. Spleens were Dounce homogenized and mononuclear cells obtained using LymphoprepTM (StemCellTechnologies). Single cell suspensions were blocked with CD16/32 (93) and stained with the following antibodies: MHCII (M5/114.15.2), CD45.2 (93), TREMl4 (16e5), CD11b (M1/70), CD115 (Afs98), CD3 (17A2), CD19 (6d5), Siglec-F (E50-2440), NK1.1 (PK136), Ly6C (hk1.4), Ly6G (1a8), CD11c (N418), CD135 (A2F10.1), Sca1 (D7), CD117 (2b8), CD127 (A7R34), TER119 (ter-119), Streptavadin (BioLegend). The cells were resuspended with DAPI, or propidium iodide (PI) solution (eBioscience) and analyzed with a 6-Laser Fortessa cytometer (BD Biosciences). For accurate cell number quantification, 123count eBeads™ Counting Beads (ThermoFisherScientifc, Cat No 01-1234-42) were used. The cells were sorted using FACS Aria Fusion (BD Biosciences). The flow cytometry data were analyzed with FlowJo (Three Star, Ashland OR).

For analysis, blood monocytes were defined as lineage^-^ (CD3, CD19, NK1.1), CD11b^+^, CD115^high^ and then assessed using both Ly6C and Treml4 to separate out Ly6C^high^ (Ly6C^high^/Treml4^-^), Ly6C^low^ (Ly6C^low^/TREML4^high^), and Ly6C^int^ populations (Supplementary Figure 1). Blood neutrophils were defined as lineage^-^, CD11b^+^, CD115^low^, and Ly6G^+^ (Supplementary Figure 1). BM monocytes were defined as lineage^-^ (CD3, CD19, NK1.1, Ter119, Ly6G), CD11b^+^, CD115^high^ and then assessed as Ly6C^high^, or Ly6C^low^ based on Ly6C expression alone (Supplementary Figure 2). BM neutrophils were identified as Lineage^+^ CD11b^+^ cells.

For analysis and sorting of BM Ly6C^high^ monocytes, Macrophage Dendritic Progenitor cells (MDPs) and Common Myeloid Progenitors (cMoPs), live, CD115^high^ cells were selected as lineage^-^ (CD3, CD19, NK1.1, Ter119) and Ly6G^-^. Sca1^-^, CD127^-^ (IL7rα-) cells were gated into three populations based on CD117 (cKit) and CD135 (Flt3) expression and each of these populations assessed for Ly6C and CD11b expression: CD117^low^, CD135^-^, Ly6C^high^, CD11b^+^ monocytes; CD117^high^, CD135^-^, Ly6C^high^, CD11b^-^ cMoPs; CD117^high^, CD135^+^, Ly6C^-^, CD11b^-^ MDPs (Supplementary Figure 3). Sorted MDPs and cMoPs were cultured, picked and cells Giemsa stained to validate identity.

For sorting of Lin^-^ Sca1 ^-^ c-Kit^+^ (LK), live cells were selected as lineage^-^ (CD3, CD19, NK1.1, Ter119) and Ly6G^-^. LK cells were gated as Ly6C^-^, CD11b^-^, CD117^+^, Sca1^-^ (Supplementary Figure 4). The observation of all types of colonies in CFU assays from these cells confirmed their multipotency, particularly the presence of red tinged BFU-E colonies (data not shown).

For analysis of primitive BM progenitors, myeloid progenitors were defined as Lineage^-^ (CD3, CD19, NK1.1, CD11b, Ly6G, Ly6C, Ter119) and CD127^-^ and then divided into LK and LSK as CD117^high^, and Sca1^-^ and Sca1^+^ respectively. The LSK population was divided into CD135^+^ Multipotent progenitors (Flt3^+^ MPPs) and CD135^-^ Haematopoietic Stem Progenitor Cells (HSPCs). All CD135^-^ cells were then divided into CD48^+^ cells, representative of the CD135^-^ MPPs (Flt3^-^ MPPs), and CD48^-^, CD150^+^ Haematopoietic Stem Cells (HSCs). The LK population was divided into Megakaryocyte-Erythrocyte Progenitors (MEPs) (CD34^-^, CD16/32), CMPs (CD34^+^, CD16/32^-^) and GMPs (CD34^+^, CD16/32) (Supplementary Figure 5).

For the spleen, following selection of live single cell populations, lineage^-^ (CD3, CD19, NK1.1, Ter119, Ly6G) and CD115^high^ cells were gated and then divided according to expression of CD135 and CD117 with the monocytes defined as double negative and MDPs as double positive. Monocytes were gated as CD11b^+^, Ly6C^high^ and MDPs as CD11b^-^, Ly6C^-^ (Supplementary Figure 6).

### CFU assays

Progenitor cells were sorted into 1.5ml Eppendorf tubes containing Iscove’s Modified Dulbecco’s Medium (IMDM) medium and cells regained by centrifugation at 400g and re-suspended in IMDM medium at a concentration of 600cells/300μl for MDPs or 800cells/300μl for LK cells. 300μl of cell suspension was added to 3ml aliquots of MethocultTM M3534 (STEMCELL Technologies) in Falcon™ Round-Bottom Polystyrene Tubes (Falcon), vortexed and plated in 35mm petri dish with duplicates for each sample. Plates were incubated at 37°C 5% v/v CO2 >= 95% humidity in a 15cm dish with an additional petri containing Dulbecco’s PBS (DPBS). Colonies were counted at 12-14 days. For morphology, colonies were picked and cytospun (Shandon Cytospin II) and then stained with Rapid Romanowsky Stain Solutions A, B, C, (TCS Biosciences Ltd;).

### Determination of cells in DNA synthesis

Mice were injected intraperitoneally with 100ml BrdU in DPBS (10mg/ml; Sigma Aldrich). Single cell suspensions were prepared and incubated with primary antibodies as specified. Live/dead stain was applied (Zombie Aqua^TM^ Fixable Viability Kit Biolegend, Fixable Viability Dye eFluor™ 780 eBioscience). Cells were fixed and washed with FoxP3-staining-buffer-set (eBioscience) and then incubated with DNase at 37°C for 30 mins (1mg/ml in D-PBS, Sigma Aldrich), DNase-solution 30ml DNase-stock + 960ml D-PBS + 10ml MgCl_2_). Cells were washed and stained with Anti-BrdU-antibody (Alexa Fluor® 488 anti-BrdU Antibody, EBioScience) for 30 mins at RT. For each group and in all experiments, background BrdU levels were set using a non-DNase treated control. Representative staining and gates are shown in Supplementary Figure 7A-C.

### RNA-seq of monocytes

For scRNAseq, the total population of monocytes (defined as lineage^-^ (CD3, CD19, NK1.1), CD11b^+^, CD115^high^) were sorted into low binding, conical 96 well non-skirted plates (Sigma) containing 2μl of lysis buffer (1.9ml of 0.2% v/v Triton-X l00 v/v + 0.1ml of RNasin Plus RNase inhibitor (10,000 U/ml, Promega). Smart-seq2 protocol was performed as described in Picelli et al [43]. Libraries were pooled in a 1:1 ratio and sequenced on one lane of an Illumina HiSeq2500. Raw reads were aligned onto mouse mm10 assembly with BWA. Only uniquely aligned reads were retained and quantified using htseq [44]. Reads were normalized using RSEM [45]. Clustering and differential gene expression analysis was done using Seurat package in R [46].

For deep RNA sequencing, monocytes were sorted as detailed above and RNA was isolated using 475μl TRIzol™ LS Reagent (Illumina). Poly A enriched mRNA was fragmented, in 2x Superscript III first-strand buffer with 10mM DTT (Invitrogen), by incubation at 94°C for 10 minutes, and immediately chilled on ice. Samples of 10 μL of fragmented mRNA, 0.5 μL of Random primer (Invitrogen), 0.5 μL of Oligo dT primer (Invitrogen), 0.5 μL of SUPERase-In (Ambion), 1 μL of dNTPs (10 mM) and 1 μL of DTT (10 mM) were heated at 50°C for one minute. At the end of incubation, 6 μL of water, 1 μL of DTT (100 mM), 0.1 μL Actinomycin D (2 μg/μL), 0.5 μL of Superscript III (Invitrogen) were added and incubated in a PCR machine using the following conditions: 25°C for 10 minutes, 50°C for 50 minutes, and a 4°C hold. The product was then purified with Agentcourt RNAClean XP beads (Beckman Coulter) according to the manufacturer’s instruction and eluted with 10 μL nuclease-free water. The RNA/cDNA double-stranded hybrid was added to 1.5 μL of Blue Buffer (Enzymatics), 1.1 μL of dUTP mix (10 mM dATP, dCTP, dGTP and 20 mM dUTP), 0.2 μL of RNase H (5 U/μL), 1.2 μL of water, 1 μL of DNA polymerase I (Enzymatics). The mixture was incubated at 16°C for 2 hours. The purified dsDNA underwent end repair by blunting, A-tailing and adaptor ligation barcoded adapters (NextFlex, Bio Scientific). Libraries were PCR-amplified for 9-14 cycles, size selected by gel extraction, quantified by Qubit dsDNA HS Assay Kit (Thermo Fisher Scientific) and sequenced on a NextSeq 500.

Data was mapped to custom genomes using STAR [47] with default parameters. All samples were assessed for quality using the FastQC (Babraham Bioinformatics, 2010) package with MutliQC [48]. Unique mapping rates and read depths were checked with a cut-off of 90% minimum uniquely mapped reads and >20×10^6^ total read depths unless otherwise specified. Additionally, correlation between samples and clonality were checked across all samples and all samples were visually inspected on the UCSC genome browser. Adapters were trimmed and counts were generated using HOMER available at http://homer.ucsd.edu/homer. Differential gene expression (DGE) was assessed with DESeq2 package with False discovery rate <-0.05 and betaPrior = TRUE. Data visualization was undertaken in R studio. For average expression and log fold change (LFC) visualisation, LFC shrinkage using alpegm method was used. Pathway analysis was undertaken using Metascape [49] http://metascape.org/gp/index.html). To convert mouse genes to human orthologues the R package g:Orth [50] was used with the following command line: orth=gorth(rownames(mousedf), source_organism = “mmusculus”, target_organism = “hsapiens”, region_query = F, numeric_ns = ““, mthreshold = 1, filter_na = T, df = T). As there are multiple orthologue matches, the genes with the highest variance were selected.

### Data Access

- Single Cell datasets are deposited in GEO: P02E01 - GSE183838 P02E02 - GSE184870 DEG in Supplemental excel files 1-3

### Statistical Analysis

All quantitative analyses were based on at least three sample replicates. Data are presented as means ±SD by GraphPad Prism. Independent-sample student t test were performed (SPSS). NS, not significant; *P < 0.05; **P < 0.01; ***P < 0.001.

## Results

### Tumor bearing MMTV-PyMT mice show increased numbers of monocytes

The effects of cancer on monocytes and neutrophils were determined in the blood, BM and spleen using the Polyoma Middle T mouse model of breast cancer on a BL6 genetic background. The diagnostic flow cytometry panels defined in the materials and methods were used together with counting beads to calculate the frequency of specific cell populations in the blood. To perform this study control and cancer bearing mice were bled bi-weekly from 8 weeks of age through to the age that the tumours were 24mm diameter corresponding to the end stage according to our approved animal protocol. As tumor onset is sporadic on the BL6 background we grouped the tumour bearing mice into those that had early and late stage carcinomas as defined histologically [40]. There was an 1.4-fold increase in the total cellularity of the blood in mice with tumours (Fig 1, A). This expansion was limited to the CD11b^+^ fraction, with no expansion in other circulating CD11b^-^ cells such as T or B cells (Fig 1, B). Using CD115 and Ly6G expression to distinguish monocytes (CD115^high^) from neutrophils (Ly6G^+^), we found that both monocytes and neutrophils increased in numbers even in control mice with age but this effect was greater in late cancers and accounted for most if not all the expansion of CD11b^+^ cells (Fig 1, C, D). However, the expansion of monocytes was more modest than observed for neutrophils (2.1 and 3.6-fold, respectively) and was restricted to mice bearing late-stage tumours while the increase in neutrophils was significant even in early cancers. Notably, the distribution of monocytic populations was unaltered, with equivalent percentages of Ly6C^high^, Ly6C^int^ and Ly6C^low^ monocytes in age-matched non-tumour control and tumour-bearing mice (Fig 1, E). Analysis of the BM from femurs also showed an expansion in monocytes and neutrophils (Fig1, F) suggesting that cancer affects myelopoiesis in this tissue and that this might contribute to the blood monocytosis and neutrophilia.

**Figure 1.**
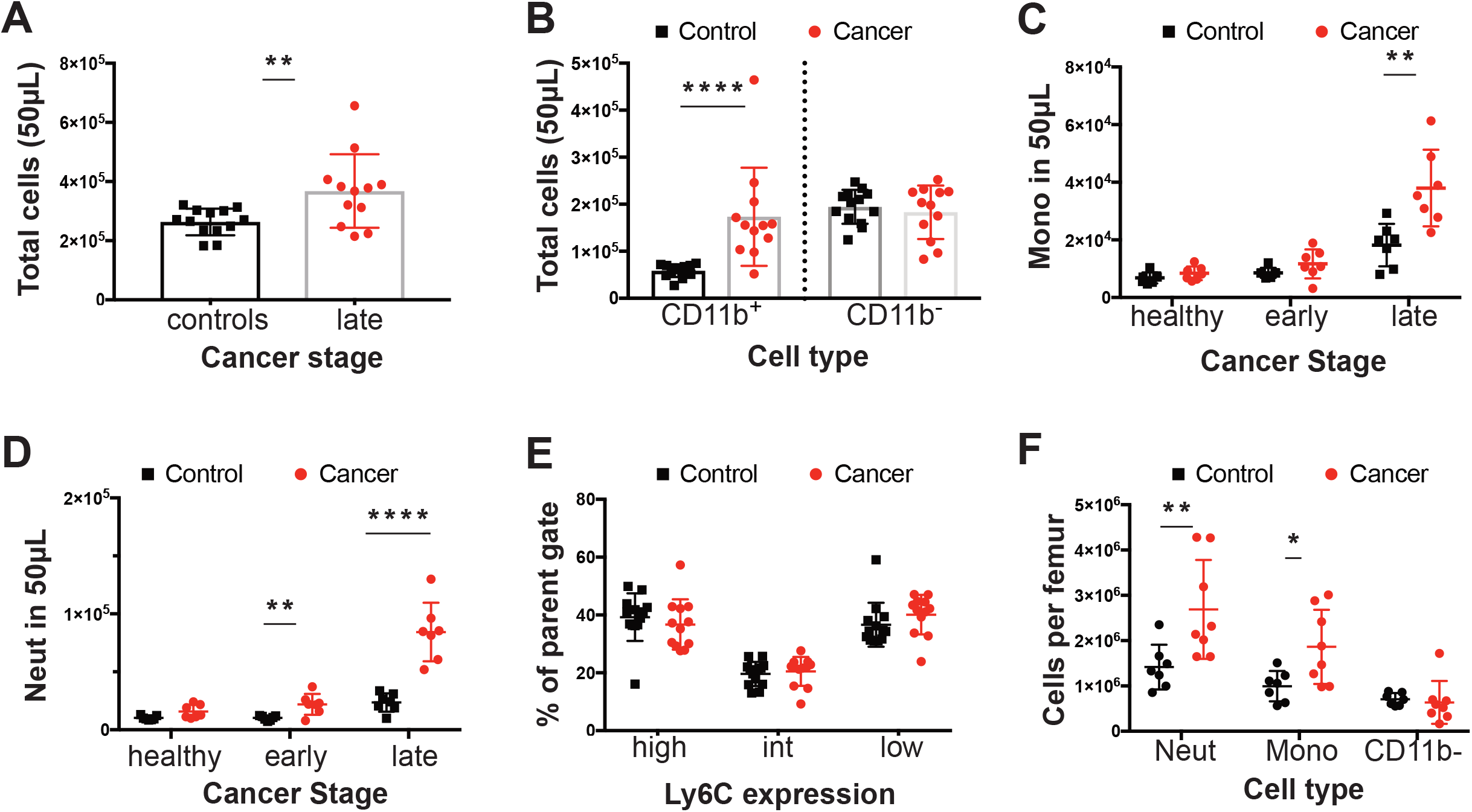
Changes in blood and bone composition of immune cells in PyMT tumour bearing mice. (A) Total cells in 50μl of blood in mice with ~20mm tumour (late) late-cancer (red dots) versus controls (black dots). (B) Total number of CD11b^+^ and CD11b^-^ cells in mice with late-stage cancer versus controls. (C) Total circulating monocytes prior to tumour development (healthy), at onset of tumours (early) and at late-stage versus control WT mice. (D) Total neutrophils (Neut) prior to tumour development (healthy), at onset of tumours (early) and at late-stage of cancer. (E) Percentage (%) of monocytes subpopulations compared at late stage with control WT mice. (F) Neutrophils, monocytes and Lineage^+^ CD11b^-^ cells per femur in mice with late-state cancer versus controls. Cancer samples are shown in red and age-matched controls in black. * p value <0.05, ** p value <0.01, ***p value <0.001, ****p value <0.0001, multiple t-test. Experiments were conducted in duplicates with littermate and co-housedgroups of n=6 (3 PyMT+ve) and n=8 (4 PyMT+ve) in each replicate for the blood and co-housed groups of n=6 (3 PyMT+ve) and n=9 (5 PyMT+ve) for the bone.

### Tumor-bearing mice exhibit elevated proliferation of BM monocytes

The purpose of this study was to study monocyte biology as neutrophil biology had been studied in detail before [42]. Thus, to examine the underlying cause of the cancer-related BM and blood monocytosis, the frequency and differentiation potential of monocyte progenitors in the BM was assessed. In the B6 PyMT model there was no increase in the frequency of either MDPs or cMoPs in the BM of tumour bearing mice. Furthermore, there was no change in the MDP upstream progenitor defined by Lin^-^, cKit^+^, Sca1^-^ cells (LK) (Figure 2, A). To support this conclusion, there was no difference in the ratio of colonies formed when LK cells were cultured in vitro in standard colony forming unit assays (Figure 2, B), nor any increase in the potential of LK, MDPs or cMoPs from mice bearing tumours to form colonies in this assay (Figure 2,C). These data suggest that both the fate potential and proliferative capacity of monocyte progenitors in the BM remain unaltered in tumour-bearing mice.

**Fig 2.**
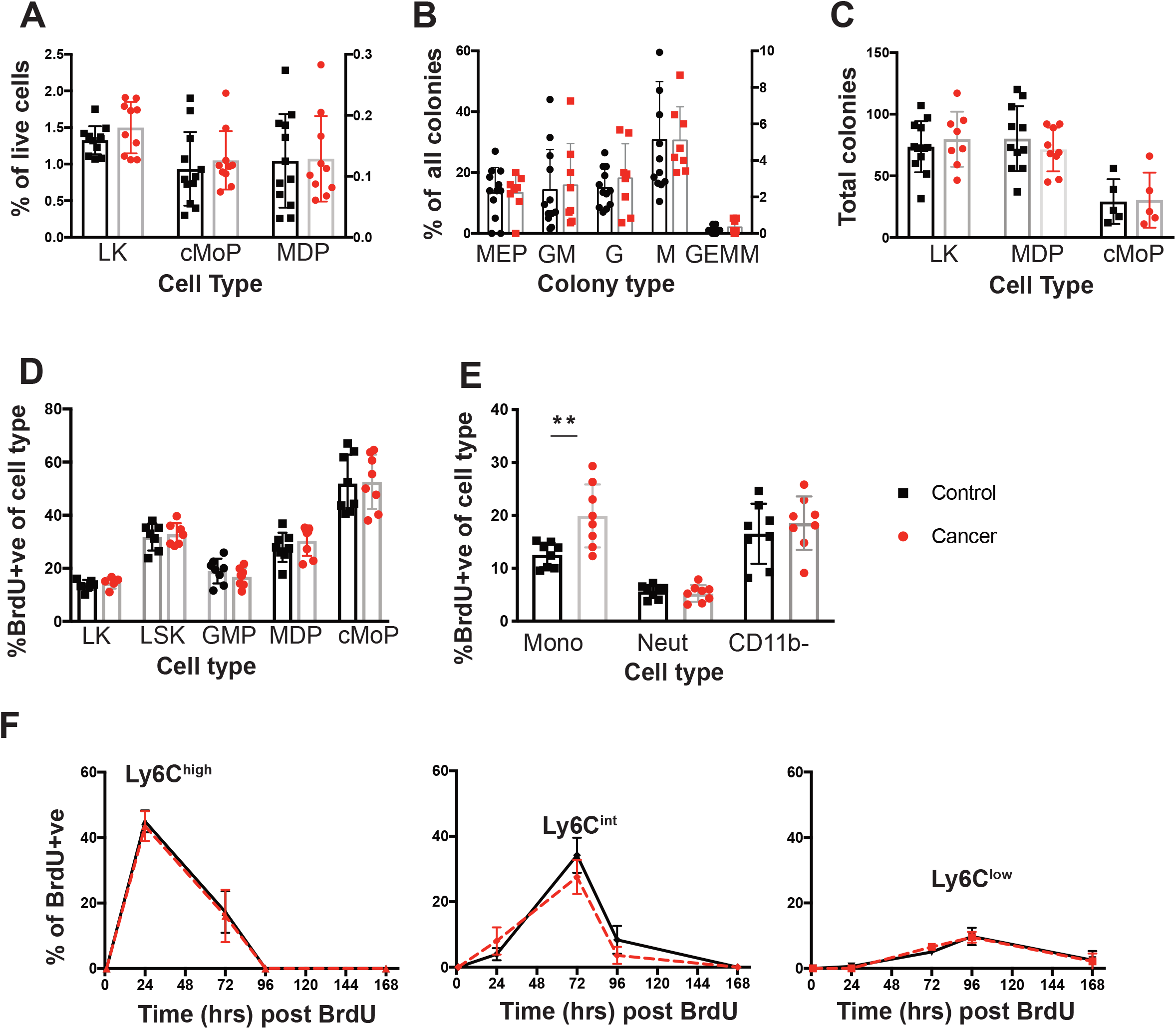
Dynamics of monopoiesis in tumor bearing mice. (A) Proportion of LK, cMoP and MDP cells within the BM shown as a percentage (%)of live cells. (B) Distribution of colony types within the LK CFU methylcellulose assays. (C) Isolated LK cells, MDPs, cMoPs progenitors were cultured in CFU assays in methocellulose for 12-14 days and the total number of colonies present were quantified. (D) Nuclear BrdU incorporation in the BM progenitors (LK, LSK, GMP, MDPs, cMoPs) 1hr post injection in mice with late cancer or age-matched controls. (E) BrdU nuclear incorporation in BM monocytes, neutrophils (Neut) and Lineage^+^ CD11b^-^ cells 1hr post injection in mice with late cancer or age-matched controls. (F) Level of BrdU^+^ circulating monocytes within each population over time. From left to right: Ly6C^high^ Ly6C^int^, Ly6C^low^. Late cancer samples are shown in red and age-matched controls in black. **p value <0.01, unpairedt-test. For LK cells sorts conducted on 3 days in triplicates of co-housed littermate groups of n=4 (2 cancer), n=11 (4 cancer) and n=5 (2 cancer). For MDPs sorts conducted on 3 days with co-housed littermate groups of n=6 (3 cancer), n=4 (2 cancer) and n=10 (5 cancer). For cMoPs sorts were conducted on 1 day with n=10 (5 cancer). For the Brd UBM experiments were conducted in triplicates of co-housed littermate groups of n=6 (3 cancer), n=6 (3 cancer) and n=4 (2 cancer). For the BrdU blood tracing, experiments were conducted in duplicates of n=6 (3 cancer, 3 controls/group).

To determine if the proliferation of these progenitors in vivo was altered by presence of cancer, we identified cells in S-phase by intraperitoneal injection of the thymidine analogue, BrdU, and analysing nuclear incorporation after 1hr. Supporting the CFU assays, there was no difference in the incorporation of BrdU between cancer and control in the Lin^-^, Sca1^+^, c-Kit^+^ (LSK), LK, GMPs, MDPs or the cMoPs (Figure 2, D). Monocytes can also proliferate in the BM in normal and cancer bearing mice [11, 19] and therefore their frequency in S-phase in vivo was also measured in the same 1 hr BrdU pulse experiment. Strikingly, mice bearing late-stage tumours exhibited significantly greater proliferation of BM monocytes, while proliferation of BM neutrophils and CD11b^-^ cells was equivalent to that found in control animals (Figure 2E).

The increased blood monocyte population in the blood could be due to either this increased proliferation in the BM and/or to their proliferation, survival and clearance once they have entered the blood. To determine the latter, pulse-labelling with BrdU was used in mice with late-stage tumours or age-matched controls to determine the half-life of blood monocytes over a 10-day period as described by [11]. No BrdU^+^ monocytes were detected in the blood at 1hr post-BrdU pulse confirming that proliferation of blood monocytes neither occurs in control nor cancer-bearing mice (Figure 2, F). Peak labelling with BrdU occurred at 24, 72, and 96 hrs for Ly6C^high^, Ly6C^int^ and Ly6C^low^ monocytes respectively (Figure 2, F), consistent with previous estimates of the sequential differentiations and different half-lives of these populations [11]. However, at no timepoint was there a difference in the level of BrdU^+^ cells apparent in the monocyte subsets from control and cancer-bearing mice (Figure 2, F). Together these findings strongly suggest that the increase in blood monocytes found in tumour-bearing mice arises from elevated proliferation of Ly6C^high^ monocytes in the BM while the survival, half-life and differentiation of these cells subsequently remains unaltered following their entry into the blood.

The spleen contains a significant reservoir of Ly6C^high^ monocytes during homeostasis [51] but may also be a site of significant extramedullary monopoiesis under conditions of stress [52, 53]. Notably, mice with late-stage cancer exhibited splenomegaly (Figure 3, A). While there was an increase in the number of mononuclear cells (MCs) in tumour bearing mice versus age-matched controls (T cells, B cells and Monocytes isolate from gradient separation) (Figure 3, B), the density of mononuclear cells per gram of spleen was equivalent in age-matched controls versus tumour bearing mice (Figure 3, C). However, splenic Ly6C^high^ monocytes formed a greater proportion of live cells and their estimated frequency was also significantly increased in tumour bearing versus age-matched controls (Figure 3, D-E). To determine if monopoiesis in the spleen contributes towards the increase in circulating monocytes observed, we looked for evidence of splenic MDPs by flow cytometry and local proliferation of monocytes at 1hr post BrdU injection. However, the frequency of splenic Ly6C^high^ monocytes that incorporated BrdU was low (1-2% BrdU^+^) in both control and tumour-bearing mice, despite proliferation of other mononuclear cells (Figure 3, F), and splenic MDPs could not be detected in control mice or tumor bearing mice (data not shown). These data suggest recruitment to the spleen from the blood is the predominant cause of the elevated monocyte count.

**Figure 3.**
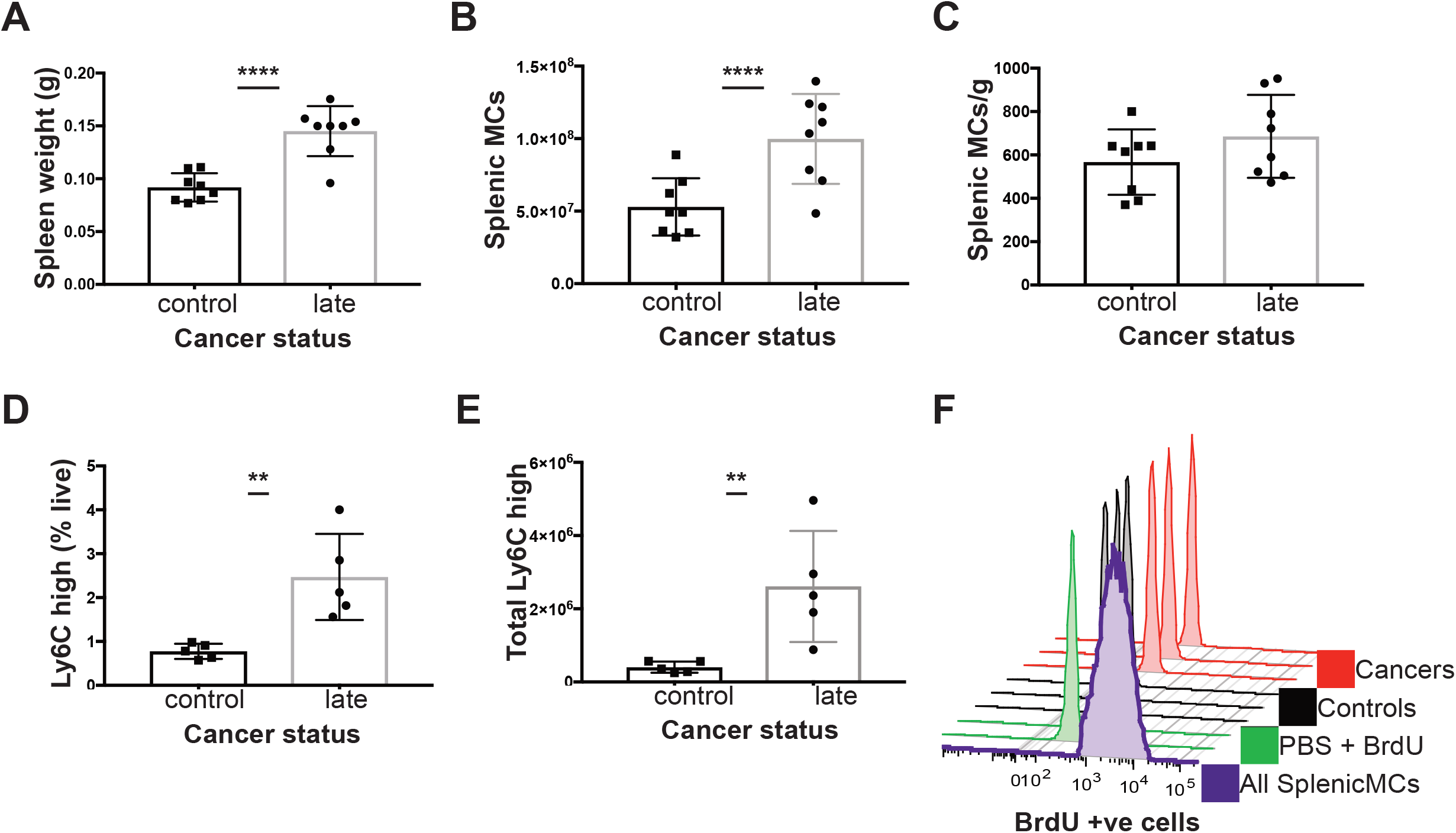
Characterisation of the spleen in C57BL/6 mice with late cancer. (A) Splenic weight. (B) Total splenic mononuclear cells (MCs; T cells, B cells, monocytes) (C) Total splenic MCs (x10^6^) normalised to splenic weight. (D) Ly6C^high^ monocytes as % of live cells. (E) Total Ly6C^high^ monocytes in spleen. (F) Histogram of splenic BrdU levels in: all Splenic MCs in spleen (purple), PBS with no DNase control (green), Ly6C^high^ Mo from controls (n=3, black) and Ly6C^high^ monocytes from late cancer mice (n=3, red) 1hr post injection of BrdU. ** p value <0.01, **** p value <0.0001, unpaired t-test. Experiments were conducted in duplicates of co-housed littermate groups of n=6 (3 cancer) and n=4 (2 cancer).

### Monocytes from the blood of mice with tumours are transcriptionally altered

Having observed a difference in the proliferation of BM monocytes in tumour bearing mice, we next examined whether cancer led to a transcriptional shift in the monocytes. First, to establish if there were transcriptionally distinct subpopulations of monocytes that differed in tumour bearing mice, single cell sequencing was performed using SMARTseq on the total monocyte population as described in the Materials and Methods. Plotting the data using a Uniform Manifold Approximation and Projection (UMAP) identified two major populations and one minor one (Figure 4Ai) and indicated that there were no new populations of monocytes occurring in response to cancer (Figure 4,A ii). Marker analysis showed the two major populations corresponded to classical (Ly6C^+^, CCR2^+^) and non-classical monocytes (Treml4^+^, NR4A1^+^) populations respectively (4A iii). As classical monocytes differentiate into non-classical ones the small population 2 is likely the intermediate monocyte population [11]. This contention is supported by the marker analysis (Figure 4Aiii) as the two defining markers for each major population are represented in this intermediate one.

**Figure 4.**
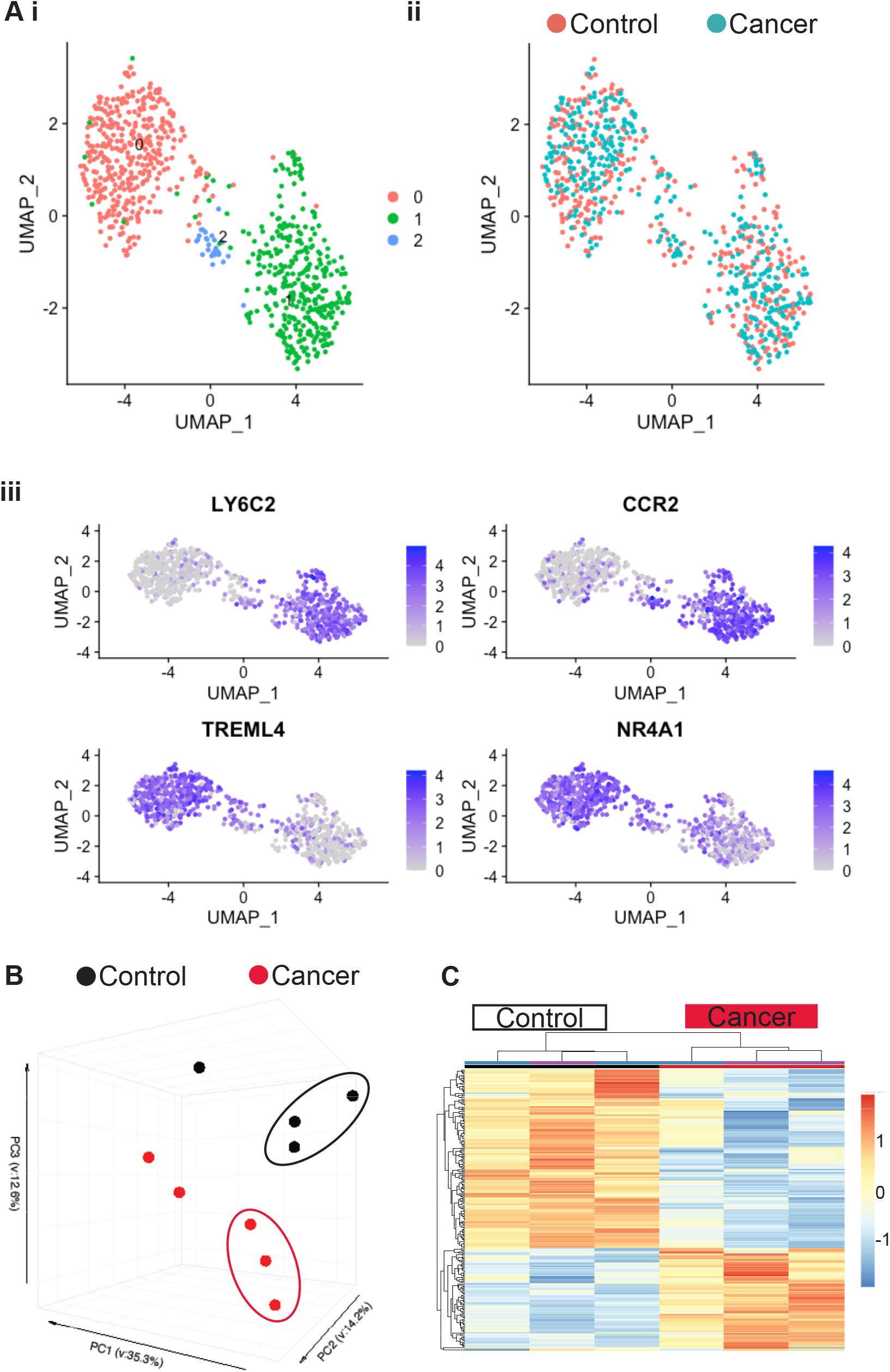
Single cell and bulk RNA-seq of blood monocytes in C57BL/6 mice with late stage cancer. (A) UMAP plot for Single cell sequencing (SCseq) of total monocytes cells from control and cancer bearing mice. i) distribution of all monocytes combined shows three populations pink, blue and green. ii) Distribution of control monocytes (pink) and tumour (blue). iii) marker analysis showing classical (LY6C2, CCR2) and non-classical (TREML4, NR4A1) monocytes. (B) PCA of gene expression derived from bulk RNAseq of Ly6C^high^ Monocytes. (C) Gene expression heatmap of Ly6C^high^ blood monocytes differentially expressed genes (DEGs) between control and cancer samples with a q.value<0.05. Samples are arranged horizontally, and sample characteristics are provided in horizontal bars for each column denoting cancer in red and control in black and the date samples were sorted in purple and blue. Genes are arranged vertically; Red colour within the heatmap indicates up-regulation, and blue colour indicates down regulation based on the tpm z-score (range [-2, 2]). Samples are clustered using complete linkage and Pearson correlation.

Having established that monocyte populations in cancer were similar to control mice, we sorted monocytes according to Ly6C expression and undertook deep RNA sequencing. Principle component analysis (PCA) revealed monocytes to primarily cluster according to monocyte subset rather than condition (Supplementary Figure 9, A). However, separate analysis of Ly6C^high^ and Ly6C^low^ populations revealed that while there was very little alteration in the Ly6C^low^ Monocytes in tumour bearing mice, (with just 7 DEGs revealed) (Supplementary Figure 8, B), the Ly6C^high^ Monocytes were altered, clustering separately from those of control mice (Figure 4, C). Ly6C^high^ Monocytes featured a total of 611 differentially expressed genes (DEGs) (adj pvalue <0.05) between cancer and control (Figure 4, D). There was an almost 2-fold difference in the number of genes down versus up-regulated (393 down, 218 up). To identify potentially important genes, the chemokines and transmembrane receptors relevant to monocyte biology were explored along with transcription factors (Figure 5, A). Key genes modulated by IFNs were downregulated. This included the type II IFN induced chemokine gene, *Cxcl10* and genes for MHCI and MHCII proteins. The lineage determining transcription factors (LDTFs), *Fosb* and *Jun,* were down-regulated. Signal transducer and activator of transcription (STATs) involved in IFN response, *Stat1* and *Stat2* were down-regulated along with the regulatory TFs *Socs1, Socs3, Socs6, Jak2* and *Jak3.* To further explore whether the DEGs may be functionally related, genes were analysed for pathway enrichment using metascape [49]. In the down-regulated list of genes, pathways were highly enriched (log q.value>10) in relation to IFN signalling and antigen presentation (Figure 5, B).

**Figure 5.**
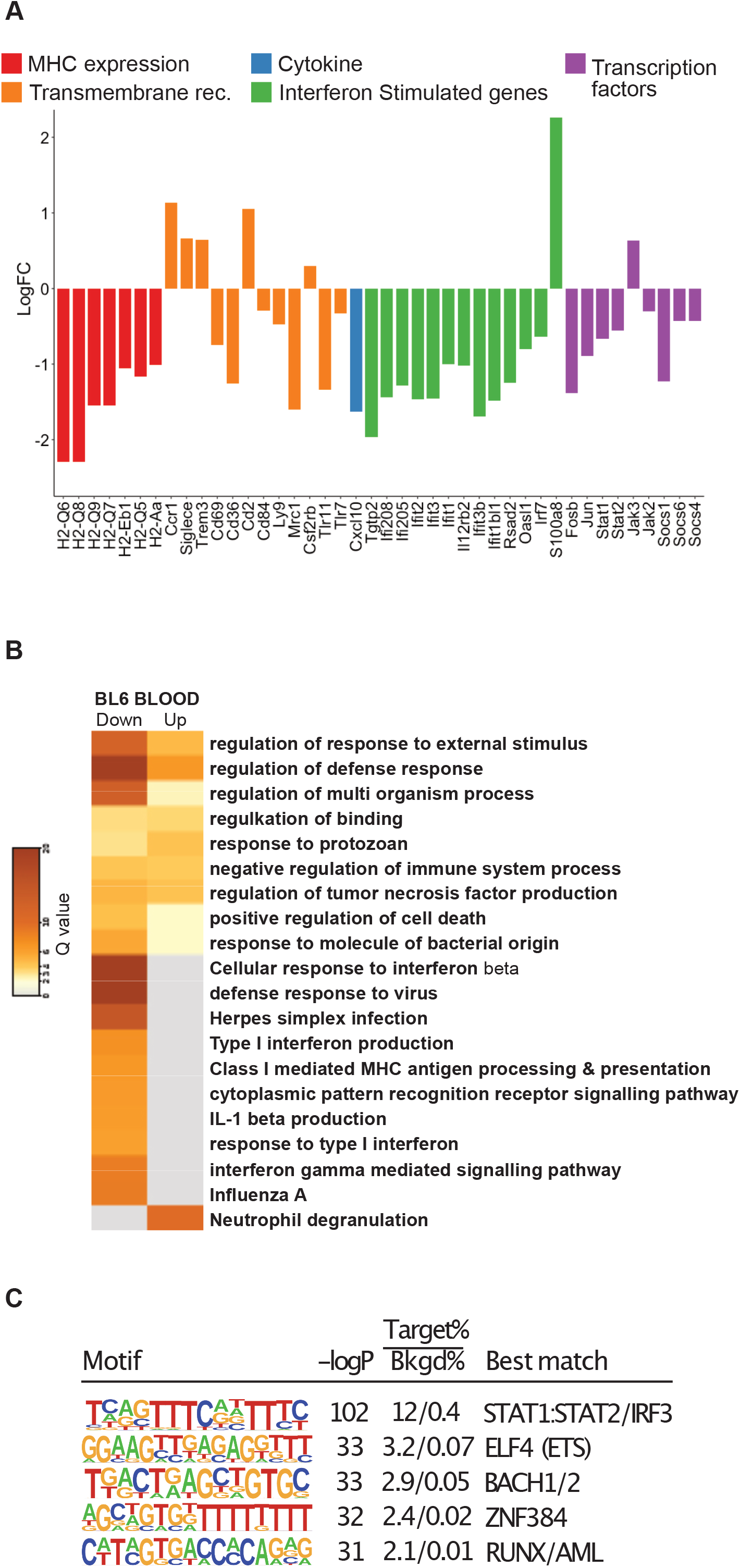
Analysis of DEGs for blood monocytes in C57BL/6 mice with late cancer. (A) Histogram of LFC in selected DEGs for blood monocytes. Genes have been selected and grouped functionally as cytokines (blue), transmembrane receptors related to MHC expression (red) and IFN response (orange), Interferon Stimulated Genes (green), and transcription factors (purple). (B) Pathways for all DEGs in the data sets using Metascape. Each column represents up or down regulated gene sets. Log q.values are plotted, and each bar coloured according to the log q.value on a scale of 0 to 30, represented with graduating intensity from cream to dark brown. Gene Ontology or KEGG terms are annotated. (C) Transcription factor *motif* enrichment analysis of the promoters of DEGs revels STAT1/STAT2 as the most highly enriched.

To investigate transcription factors associated with down-regulated genes identified in Figure 4C, we performed de novo motif enrichment analysis of their corresponding promoters. This analysis identified a motif for STAT1:STAT2 as the most highly enriched motif (Figure 5C), providing evidence that down-regulation of the mRNAs encoding STAT1 and STAT2 is functionally related to genes exhibiting reduced expression in Ly6C monocytes of tumor-bearing mice. In contrast, motifs recognized by FOSB or JUN were not highly enriched. This could reflect functional redundancy of the FOS/JUN family of transcription factors or the possibility that they primarily function at distal regulatory elements. The remaining highly enriched motifs (ELF, BACH, ZNF384 and RUNX) correspond to binding sites for factors that typically exhibit high enrichment at promoters.

### Transcriptional changes occur within the bone marrow

The above data indicates there are transcriptional alterations to circulating monocytes in the context of cancer. Hence, we wanted to ascertain at what stage of their development this conditioning may be taking place. For this purpose, the dominant Ly6C^high^ monocytic population representing ~90% of these cells from the BM were sequenced. PCA revealed that cancer samples clustered separately from controls (Figure 6, A). Further analysis revealed 158 genes to be differentially expressed between cancer and control samples (50 up and 108 down-regulated in cancer with a adj pvalue<0.05) (Figure 6, B). Even though there were fewer genes altered in BM monocytes just under half of those genes that are down-regulated in BM monocytes by cancer were also down-regulated in Ly6C^high^ blood Monocytes (Figure 6, C). These 47 common genes were highly enriched for type I IFN pathways (Figure 6, D).

**Figure 6.**
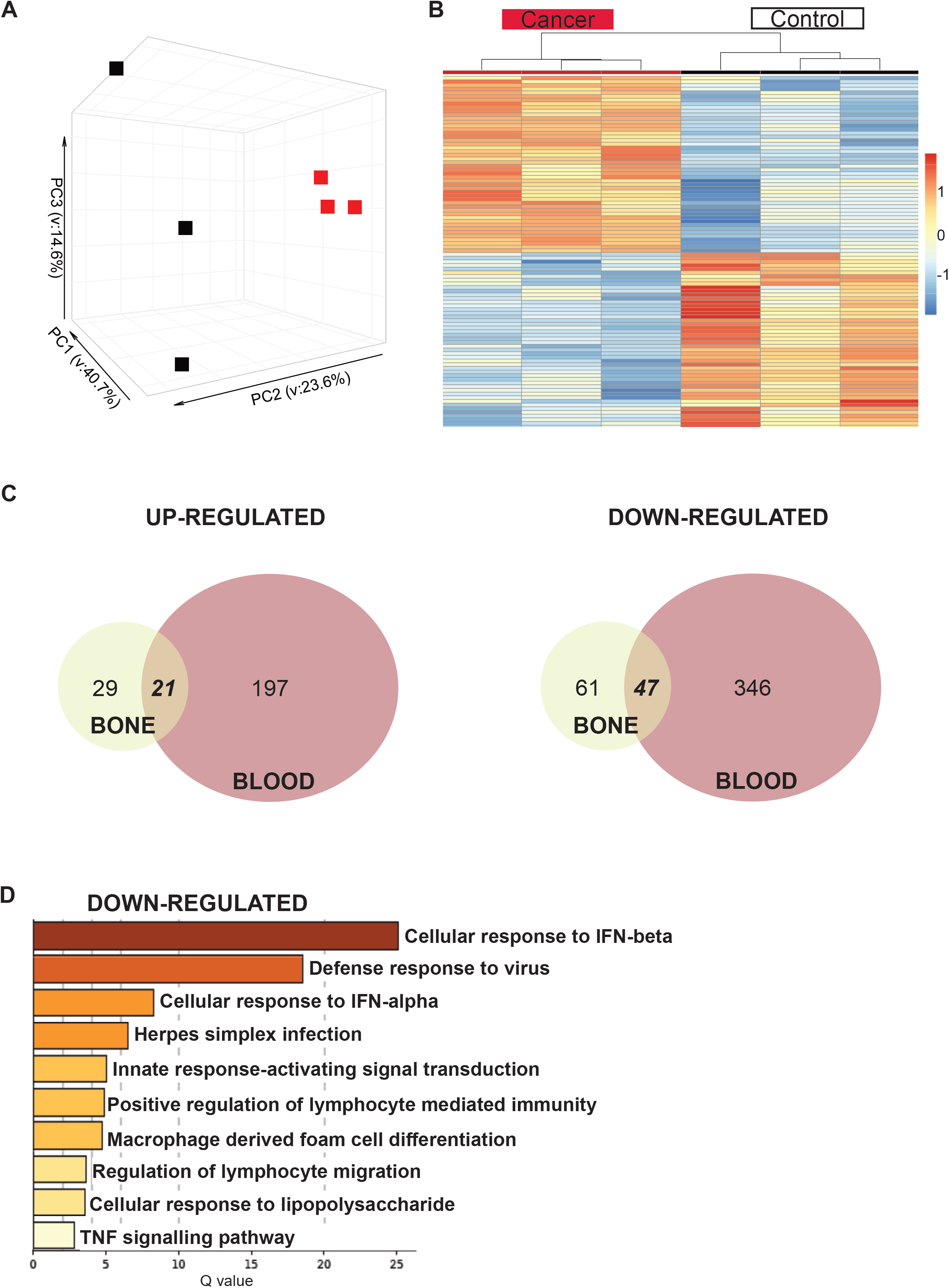
Analysis of RNA-seq of Ly6C^high^ bone and blood monocytes. (A) PCA of Ly6c^high^ bone monocytes. (B) Gene expression heatmap of Ly6C^high^ bone monocytes DEGs between control and cancer samples with a q.value<0.05. Samples are arranged horizontally, and sample characteristics are provided in horizontal bars for each column denoting cancer in red and control in black and the date samples were sorted in purple and blue. Genes are arranged vertically; Red colour within the heatmap indicates up-regulation, and blue colour indicates down regulation based on the tpm z-score (range [-2, 2]). Samples are clustered using complete linkage and Pearson correlation. (C) Venn diagram for DEGs in BM (cream) and blood (red) that are up-regulated and down-regulated in cancer. (D) Pathways for 47 shared down-regulated genes. Log q.values are plotted, and each bar coloured according to the log q.value on a scale of 0 to 30, represented with graduating intensity from cream to dark brown. Gene Ontology or KEGG terms are annotated.

### Transcriptional shifts in the mouse PyMT model are not orthologous to patients with Breast Cancer

Having identified the DEGs in cancer and control mice and the relevant pathways that were altered these were compared to the human data on total circulating monocytes (classical and non-classical) in patients with breast cancer and healthy controls [36]. To undertake this comparison a list of 865 DEGs between the breast cancer patient and control healthy volunteer samples was used (selected using a criterion of absolute LFC>1.5). Comparing these datasets very few DEGs were common to both mouse and human (Figure 7 A, B). In case the strain of the mouse may affect the findings, circulating monocytes were also sequenced from FVB mice with PyMT tumours and age-matched controls. This generated a third list of DEGs (see supplementary files S3). However, again there were very few DEGs common to mouse and human (Figure 7 A, B, C). The Lineage determining transcription factor, *Jun* was the only gene that was commonly down-regulated in all the DEG lists. To identify if there may be common pathways affected, despite a lack in shared DEGs, joint pathways analysis was undertaken. This revealed some commonly enriched down-regulated pathways, particularly with respect to metabolism, cell differentiation, survival and migration. However, the main pathway that had been identified in classical monocytes in mice, IFN signalling, was up-regulated in human cancer monocytes (Figure 7 D). These data suggest different responses to cancer in the two species.

**Figure 7.**
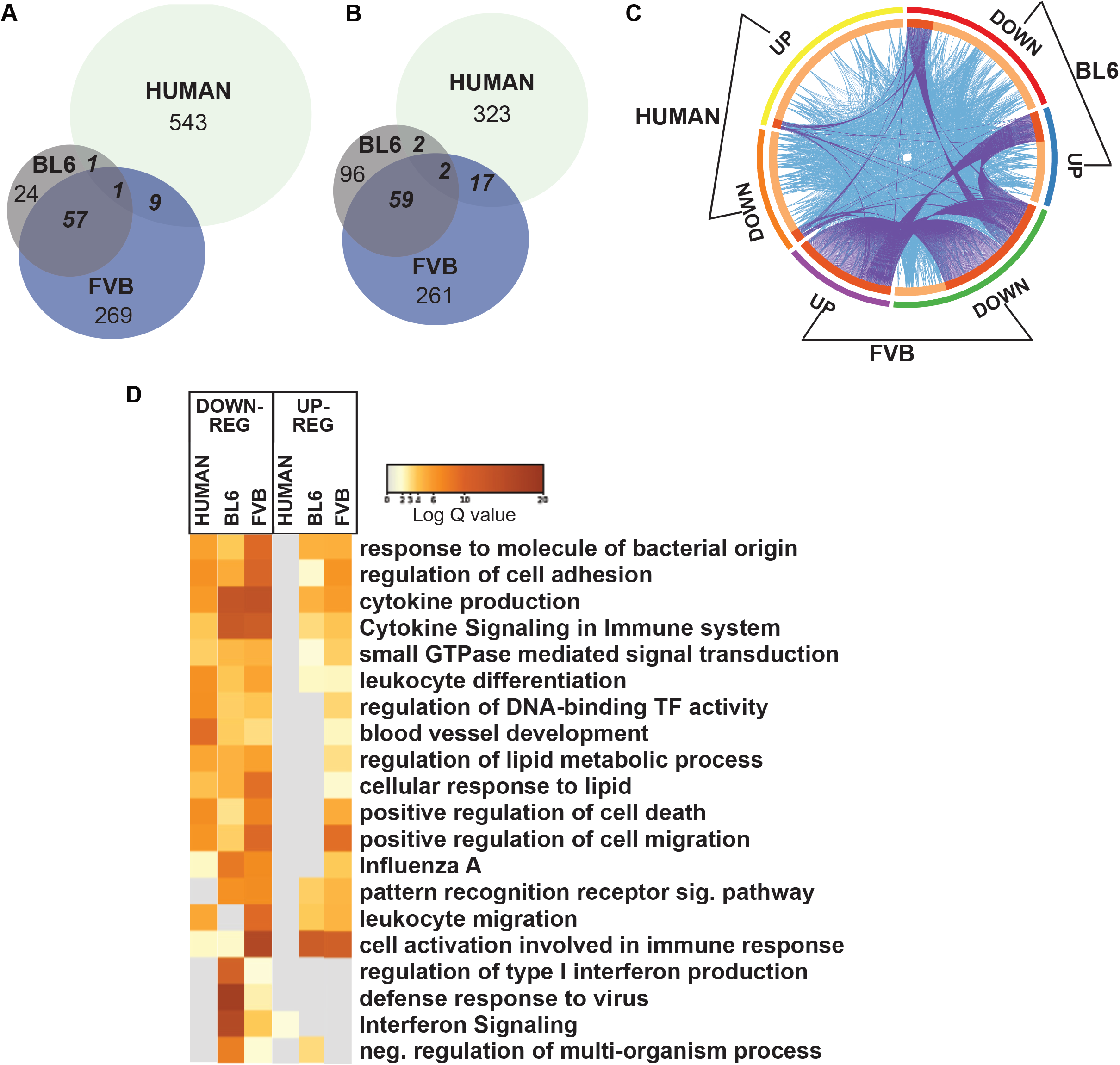
Comparison of DEGs of blood monocytes between cancer and controls samples in mouse strains (FVB and C57BL/6) and human. (A) Venn diagram of up-regulated DEGs in *Ly6C^high^ monocytes*. (B) Venn diagram of down-regulated DEGs in *Ly6C^high^ monocytes*. (C) Circos plot showing commonly enriched genes. The outer arcs represent which DEG gene list is featured. BL6; down-regulated (red arc) up-regulated (blue arc), FVB; down-regulated (green arc), up-regulated (purple arc), Human; down-regulated (orange arc), up-regulated (yellow arc). The inner arcs reflect whether the genes are common to all 3 (coloured in dark orange) or genes are common to just 2 groups (coloured in light orange). Each occasion that a gene is in common between a group a purple line occurs between where the gene occurs on the respective arcs. Each occasion a gene that belongs to the same enriched ontology term as another gene in another group a blue line occurs between where the gene occurs on the respective arcs. (D) Pathways for shared genes. Each column represents human, BL6 or FVB DEGs. Columns are grouped into pathways that were down-regulated (left) or up-regulated (right). Log q.values are plotted, and each bar coloured according to the log q.value on a scale of 0 to 20, represented with graduating intensity from cream to dark brown. Gene Ontology or KEGG terms are annotated.

## Discussion

It is well known that cancer causes systemic effects a process particularly well documented by the induction of cachexia [54]. This metabolic disease tends to occur at advanced stages of the disease. However, data is accumulating in mice and humans that cancer also causes systemic effects at early stages of the disease. An example of this phenomena is the establishment of the “pre-metastatic niche’ by early tumors that biases the homing of metastatic cells and enhances their seeding [55, 56]. The formation of this niche depends on secretion of ECM components such as fibronectin and exosomes as well as mobilization to these sites of BM derived cells such as monocytes and neutrophils [18]. The earliest descriptions of what some have described as sterile inflammation, was the accumulation of CD11b+ VEGFR1 + myeloid cells [57] that are most likely derived from classical monocytes that undergo differentiation in response to the “niche” environment [35, 58]. This pre-metastatic niche concept has yet to be established in humans, but there are increasing numbers of reports that cancer alters monocyte biology. This alteration is manifest by the enhancement monocyte number [59] and an increase in the non-classical monocyte population [13, 37] and alterations in total monocyte transcriptomes [13, 38, 39]. In two of these cases, transcriptional signatures were derived in breast and colorectal cancer that were able to predict the presence of cancer [13, 39]. However, these signatures did not overlap perhaps due to different cancers or to different transcriptional profiling methodologies.

In this study, we hypothesized that mouse models of breast cancer would have similar impacts on monocytes as found in humans and thereby allow experimental manipulation to determine mechanism. However, whilst we documented a significant monopoiesis there were no changes in non-classical to classical monocyte ratios in the blood nor any indications of new populations by FACS or Single cell RNAseq. This failure to see ratio changes in mice may simply reflect starting populations of classical to non-classical monocytes which in humans are approximately 9 to 1 but in mice these populations are approximately equal. Furthermore, there were no other significant monocytic populations such as the myeloid derived suppressor cells indicated by the analysis of side-scatter as also found in the blood of human cancers [60] nor emergence of other monocytic subtypes such as the SatM monocytes found during fibrosis [61]. The failure to find such populations might be related to the sorting procedure that was directed by traditional monocyte markers.

Our data also indicated that the monocytopoiesis induced by cancer was not due to increase in BM progenitors nor to changed half-lives of circulating monocytes but to enhanced proliferation of monocytes in the BM. Others have shown the spleen as a source of monocyte progenitors in lung cancer models [62] as a result of angiotensin II produced by cancer remotely increasing HSC frequency that migrate to the spleen [63]. However, although the number of splenic monocytes increased along with the splenomegaly, we found no evidence that cancer increased their proliferation nor enhanced the very low frequency of progenitors in in the spleen in this model of breast cancer.

Elevation of CSF1 is perhaps the most obvious explanation for increased proliferation of monocytes in the BM. Elevated levels of circulating CSF1, a proliferation and survival factor for monocytes and macrophages, at late stages of cancer in the spontaneous FVB PyMT model of breast cancer have been reported [42] as well as in human cancers [64, 65]. In the PYMT mouse model this elevation was preceded by elevated G-CSF *(Csf3)* and an earlier expansion of neutrophils when compared to monocytes, paralleling our findings [42]. Similarly, in the circulation the lack of changes in half-life of increased numbers of monocytes in cancer suggest the presence of sufficient CSF1 to maintain their viability. In the BM Interferon signalling has also been shown to inhibit proliferation of differentiated hematopoietic cells, but not to inhibit HSC proliferation [66]. Thus down-regulation of this signalling pathway as observed in our sequencing data combined with elevated CSF1 could explain the relatively specific effect of tumors on BM proliferation of monocytes.

Transcriptomic analysis indicated relatively small changes in the classical monocyte population in response to cancer. However, the translation of these findings to human is limited by the lack of orthologous genes. In our data, the greatest variance was according to monocyte subset. The human dataset that was used for comparison was collected using the entire monocytic population. The proportion of non-classical monocytes in patients with breast cancer has been shown to be higher than in healthy controls [13] and this may have masked more subtle effects of cancer on the classical population. Single cell RNAseq data comparing lung tissue isolated from Murine KP1.9 lung adenocarcinoma and 7 human primary NSCLC sample suggest that there should be homologous genes between human and mouse [8]. In mouse models, the immune profile of the tumor microenvironment is known to be specific to the genetic mutations within the tumour and the same is thought to apply to human cancers [29]. The systemic influences of cancer are also dependent on specific gene mutations [29]. The PyMT model is beneficial because of robust cancer development with histology that mirrors human ductal adenocarcinoma and metastatic capacity. However, this model is driven by the expression of a viral oncoprotein [40] thus the mutational profile does not reflect those found in human breast tumours. Furthermore, even when the cancer is driven by the same oncoprotein there were significant differences in gene expression according to strain (FVB vs BL6). In addition, the development of mouse mammary cancer in this model is over a few weeks while in humans it can be over many years. These differences limit exact comparisons between the species but do suggest mouse models maybe inadequate for exploring mechanisms behind systemic changes seen in humans.

Nonetheless, the findings in mice are still interesting. The proliferation of Ly6C^hi^ BM monocytes leads to elevated number of blood monocytes, that then circulate with normal half-life. Similar increases in monocyte number have been described in human cancers. Comparison of differentially expressed genes in the BM samples with the larger numbers of genes differentially expressed in the blood compared to BM suggest that the major systemic influence of cancer occurs in the circulation. Nonetheless, the major pathways involving interferon signalling were also changed in the bone. Overall, this suggests a two-step process whereby cancer affects transcription and proliferation in Ly6C^hi^ monocytes in the bone and once monocytes enter the circulation, further and more significant transcriptional differences are overlaid upon pre-existing changes.

## Supplementary Materials

*S1_MOUSE_ C57BL/6_BLOOD_DEGs.xls*

*S2_MOUSE_ C57BL/6_BM.xls*

*S3_MOUSE(FVB)_BLOOD_DEGs.xls*

## Acknowledgments

We would like to acknowledge Will Mungal who helped with experimental procedures. The research was funded CRUK C17950/A26783, Wellcome Trust (101067/Z/13/Z) and MRC Centre grant MR/N022556/1 to JWP and CRUK Clinical Research Fellowship (C157/A20919) to AR. MRC grant MR/L008076/1 to SJJ. NIH grant DK091183 to CKG. C.Z.H. was supported by the Cancer Research Institute Irvington Postdoctoral Fellowship Program

## Author Contributions

A C Robinson Conceptualization Data curation; Investigation: Formal analysis; writing original draft.

P Louwe Data curation; Formal analysis

S.Webb Investigation: Data curation; Formal analysis; Project administration

M Burgess Investigation; Formal analysis.

C Han Investigation; Formal analysis.

D Skola Data curation; Formal analysis

Z Ouyang Data curation; Formal analysis.

N Batada Investigation; Data curation; Formal analysis.

VG Huici, Formal analysis

L Cassetta, Data curation; Formal analysis;

C Glass Conceptualisation; Methodology; Supervision, Funding acquisition.

SJ Jenkins Conceptualisation, Methodology; Supervision: Review of the manuscript.

JW Pollard Conceptualisation; Visualization: Supervision: Methodology, Formal analysis; Writing original and final manuscripts. Project administration, Funding acquisition

## References

[1] F. Geissmann, S. Jung, D.R. Littman, Blood monocytes consist of two principal subsets with distinct migratory properties, Immunity, 19 (2003) 71–82.

[2] J. Hettinger, D.M. Richards, J. Hansson, M.M. Barra, A.C. Joschko, J. Krijgsveld, M. Feuerer, Origin of monocytes and macrophages in a committed progenitor, Nature immunology, 14 (2013) 821–830.

[3] Z. Liu, Y. Gu, S. Chakarov, C. Bleriot, I. Kwok, X. Chen, A. Shin, W. Huang, R.J. Dress, C.A. Dutertre, A. Schlitzer, J. Chen, L.G. Ng, H. Wang, Z. Liu, B. Su, F. Ginhoux, Fate Mapping via Ms4a3-Expression History Traces Monocyte-Derived Cells, Cell, 178 (2019) 1509–1525 e1519.

[4] A. Yanez, S.G. Coetzee, A. Olsson, D.E. Muench, B.P. Berman, D.J. Hazelett, N. Salomonis, H.L. Grimes, H.S. Goodridge, Granulocyte-Monocyte Progenitors and Monocyte-Dendritic Cell Progenitors Independently Produce Functionally Distinct Monocytes, Immunity, 47 (2017) 890–902 e894.

[5] R. van Furth, Z.A. Cohn, J.G. Hirsch, J.H. Humphrey, W.G. Spector, H.L. Langevoort, The mononuclear phagocyte system: a new classification of macrophages, monocytes, and their precursor cells, Bull. World Health Organ., 46 (1972) 845–852.

[6] N.V. Serbina, E.G. Pamer, Monocyte emigration from bone marrow during bacterial infection requires signals mediated by chemokine receptor CCR2, Nature immunology, 7 (2006) 311–317.

[7] M.A. Ingersoll, R. Spanbroek, C. Lottaz, E.L. Gautier, M. Frankenberger, R. Hoffmann, R. Lang, M. Haniffa, M. Collin, F. Tacke, A.J. Habenicht, L. Ziegler-Heitbrock, G.J. Randolph, Comparison of gene expression profiles between human and mouse monocyte subsets, Blood, 115 (2010) e10–19.

[8] R. Zilionis, C. Engblom, C. Pfirschke, V. Savova, D. Zemmour, H.D. Saatcioglu, I. Krishnan, G. Maroni, C.V. Meyerovitz, C.M. Kerwin, S. Choi, W.G. Richards, A. De Rienzo, D.G. Tenen, R. Bueno, E. Levantini, M.J. Pittet, A.M. Klein, Single-Cell Transcriptomics of Human and Mouse Lung Cancers Reveals Conserved Myeloid Populations across Individuals and Species, Immunity, 50 (2019) 1317–1334 e1310.

[9] A.M. Zawada, K.S. Rogacev, B. Rotter, P. Winter, R.R. Marell, D. Fliser, G.H. Heine, SuperSAGE evidence for CD14++CD16+ monocytes as a third monocyte subset, Blood, 118 (2011) e50–61.

[10] A.C. Villani, R. Satija, G. Reynolds, S. Sarkizova, K. Shekhar, J. Fletcher, M. Griesbeck, A. Butler, S. Zheng, S. Lazo, L. Jardine, D. Dixon, E. Stephenson, E. Nilsson, I. Grundberg, D. McDonald, A. Filby, W. Li, P.L. De Jager, O. Rozenblatt-Rosen, A.A. Lane, M. Haniffa, A. Regev, N. Hacohen, Single-cell RNA-seq reveals new types of human blood dendritic cells, monocytes, and progenitors, Science, 356 (2017).

[11] S. Yona, K.W. Kim, Y. Wolf, A. Mildner, D. Varol, M. Breker, D. Strauss-Ayali, S. Viukov, M. Guilliams, A. Misharin, D.A. Hume, H. Perlman, B. Malissen, E. Zelzer, S. Jung, Fate mapping reveals origins and dynamics of monocytes and tissue macrophages under homeostasis, Immunity, 38 (2013) 79–91.

[12] A.A. Patel, Y. Zhang, J.N. Fullerton, L. Boelen, A. Rongvaux, A.A. Maini, V. Bigley, R.A. Flavell, D.W. Gilroy, B. Asquith, D. Macallan, S. Yona, The fate and lifespan of human monocyte subsets in steady state and systemic inflammation, J. Exp. Med., 214 (2017) 1913–1923.

[13] L. Cassetta, S. Fragkogianni, A.H. Sims, A. Swierczak, L.M. Forrester, H. Zhang, D.Y.H. Soong, T. Cotechini, P. Anur, E.Y. Lin, A. Fidanza, M. Lopez-Yrigoyen, M.R. Millar, A. Urman, Z. Ai, P.T. Spellman, E.S. Hwang, J.M. Dixon, L. Wiechmann, L.M. Coussens, H.O. Smith, J.W. Pollard, Human Tumor-Associated Macrophage and Monocyte Transcriptional Landscapes Reveal Cancer-Specific Reprogramming, Biomarkers, and Therapeutic Targets, Cancer cell, 35 (2019) 588–602 e510.

[14] J.A. Joyce, J.W. Pollard, Microenvironmental regulation of metastasis, Nature reviews. Cancer, 9 (2009) 239–252.

[15] B.K. Stansfield, D.A. Ingram, Clinical significance of monocyte heterogeneity, Clin Transl Med, 4 (2015) 5.

[16] L. Shi, Z. Zhang, L. Song, Y.T. Leung, M.A. Petri, K.E. Sullivan, Monocyte enhancers are highly altered in systemic lupus erythematosus, Epigenomics, 7 (2015) 921–935.

[17] C. Bergenfelz, A.M. Larsson, K. von Stedingk, S. Gruvberger-Saal, K. Aaltonen, S. Jansson, H. Jernstrom, H. Janols, M. Wullt, A. Bredberg, L. Ryden, K. Leandersson, Systemic Monocytic-MDSCs Are Generated from Monocytes and Correlate with Disease Progression in Breast Cancer Patients, PloS one, 10 (2015) e0127028.

[18] E. Guc, J.W. Pollard, Redefining macrophage and neutrophil biology in the metastatic cascade, Immunity, 54 (2021) 885–902.

[19] E.N. Arwert, A.S. Harney, D. Entenberg, Y. Wang, E. Sahai, J.W. Pollard, J.S. Condeelis, A Unidirectional Transition from Migratory to Perivascular Macrophage Is Required for Tumor Cell Intravasation, Cell reports, 23 (2018) 1239–1248.

[20] K. Movahedi, D. Laoui, C. Gysemans, M. Baeten, G. Stange, J. Van den Bossche, M. Mack, D. Pipeleers, P. In’t Veld, P. De Baetselier, J.A. Van Ginderachter, Different tumor microenvironments contain functionally distinct subsets of macrophages derived from Ly6C(high) monocytes, Cancer Res., 70 (2010) 5728–5739.

[21] R.A. Franklin, W. Liao, A. Sarkar, M.V. Kim, M.R. Bivona, K. Liu, E.G. Pamer, M.O. Li, The Cellular and Molecular Origin of Tumor-Associated Macrophages, Science, (2014).

[22] B.Z. Qian, Y. Deng, J.H. Im, R.J. Muschel, Y. Zou, J. Li, R.A. Lang, J.W. Pollard, A distinct macrophage population mediates metastatic breast cancer cell extravasation, establishment and growth, PloS one, 4 (2009) e6562.

[23] B.Z. Qian, J. Li, H. Zhang, T. Kitamura, J. Zhang, L.R. Campion, E.A. Kaiser, L.A. Snyder, J.W. Pollard, CCL2 recruits inflammatory monocytes to facilitate breast-tumour metastasis, Nature, 475 (2011) 222–225.

[24] B.Z. Qian, J.W. Pollard, Macrophage diversity enhances tumor progression and metastasis, Cell, 141 (2010) 39–51.

[25] R. Noy, J.W. Pollard, Tumor-associated macrophages: from mechanisms to therapy, Immunity, 41 (2014) 49–61.

[26] T. Kitamura, B.Z. Qian, J.W. Pollard, Immune cell promotion of metastasis, Nature reviews. Immunology, 15 (2015) 73–86.

[27] Y. Zhu, M. Herndon, J., D. Sojka, K., K. Kim, B. Knolhoff, L., C. Zho, D. Cullinan, R., J. Luo, A. Bearden, R., K. Lavine, J., W. Yokoyama, M., W. Hawkins, G., R. Fields, C,, G. Randolph, J, D. DeNardo, G., Tissue Resident Macrophages in Pancreatic Ductal Adenocarcinoma Originate from Embryonic Hematopoiesis and Promote Tumor Progression, Immunity, 47 (2017) 323–338.

[28] E. Friebel, K. Kapolou, S. Unger, N.G. Nunez, S. Utz, E.J. Rushing, L. Regli, M. Weller, M. Greter, S. Tugues, M.C. Neidert, B. Becher, Single-Cell Mapping of Human Brain Cancer Reveals Tumor-Specific Instruction of Tissue-Invading Leukocytes, Cell, 181 (2020) 1626–1642 e1620.

[29] M.D. Wellenstein, K.E. de Visser, Cancer-Cell-Intrinsic Mechanisms Shaping the Tumor Immune Landscape, Immunity, 48 (2018) 399–416.

[30] R.N. Kaplan, R.D. Riba, S. Zacharoulis, A.H. Bramley, L. Vincent, C. Costa, D.D. MacDonald, D.K. Jin, K. Shido, S.A. Kerns, Z. Zhu, D. Hicklin, Y. Wu, J.L. Port, N. Altorki, E.R. Port, D. Ruggero, S.V. Shmelkov, K.K. Jensen, S. Rafii, D. Lyden, VEGFR1-positive haematopoietic bone marrow progenitors initiate the pre-metastatic niche, Nature, 438 (2005) 820–827.

[31] S. Hiratsuka, A. Watanabe, H. Aburatani, Y. Maru, Tumour-mediated upregulation of chemoattractants and recruitment of myeloid cells predetermines lung metastasis, Nature cell biology, 8 (2006) 1369–1375.

[32] Y. Liu, X. Cao, Characteristics and Significance of the Pre-metastatic Niche, Cancer cell, 30 (2016) 668–681.

[33] T. Kitamura, D. Doughty-Shenton, L. Cassetta, S. Fragkogianni, D. Brownlie, Y. Kato, N. Carragher, J.W. Pollard, Monocytes Differentiate to Immune Suppressive Precursors of Metastasis-Associated Macrophages in Mouse Models of Metastatic Breast Cancer, Frontiers in immunology, 8 (2017) 2004.

[34] T. Kitamura, B.Z. Qian, D. Soong, L. Cassetta, R. Noy, G. Sugano, Y. Kato, J. Li, J.W. Pollard, CCL2-induced chemokine cascade promotes breast cancer metastasis by enhancing retention of metastasis-associated macrophages, J. Exp. Med., 212 (2015) 1043–1059.

[35] A.M. Gil-Bernabe, S. Ferjancic, M. Tlalka, L. Zhao, P.D. Allen, J.H. Im, K. Watson, S.A. Hill, A. Amirkhosravi, J.L. Francis, J.W. Pollard, W. Ruf, R.J. Muschel, Recruitment of monocytes/macrophages by tissue factor-mediated coagulation is essential for metastatic cell survival and premetastatic niche establishment in mice, Blood, 119 (2012) 3164–3175.

[36] L. Cassetta, E.S. Baekkevold, S. Brandau, A. Bujko, M.A. Cassatella, A. Dorhoi, C. Krieg, A. Lin, K. Lore, O. Marini, J.W. Pollard, M. Roussel, P. Scapini, V. Umansky, G.J. Adema, Deciphering myeloid-derived suppressor cells: isolation and markers in humans, mice and non-human primates, Cancer Immunol. Immunother., 68 (2019) 687–697.

[37] A.L. Feng, J.K. Zhu, J.T. Sun, M.X. Yang, M.R. Neckenig, X.W. Wang, Q.Q. Shao, B.F. Song, Q.F. Yang, B.H. Kong, X. Qu, CD16+ monocytes in breast cancer patients: expanded by monocyte chemoattractant protein-1 and may be useful for early diagnosis, Clin. Exp. Immunol., 164 (2011) 57–65.

[38] M. Chittezhath, M.K. Dhillon, J.Y. Lim, D. Laoui, I.N. Shalova, Y.L. Teo, J. Chen, R. Kamaraj, L. Raman, J. Lum, T.P. Thamboo, E. Chiong, F. Zolezzi, H. Yang, J.A. Van Ginderachter, M. Poidinger, A.S. Wong, S.K. Biswas, Molecular profiling reveals a tumor-promoting phenotype of monocytes and macrophages in human cancer progression, Immunity, 41 (2014) 815–829.

[39] A. Hamm, H. Prenen, W. Van Delm, M. Di Matteo, M. Wenes, E. Delamarre, T. Schmidt, J. Weitz, R. Sarmiento, A. Dezi, G. Gasparini, F. Rothe, R. Schmitz, A. D’Hoore, H. Iserentant, A. Hendlisz, M. Mazzone, Tumour-educated circulating monocytes are powerful candidate biomarkers for diagnosis and disease follow-up of colorectal cancer, Gut, 65 (2016) 990–1000.

[40] E.Y. Lin, J.G. Jones, P. Li, L. Zhu, K.D. Whitney, W.J. Muller, J.W. Pollard, Progression to malignancy in the polyoma middle T oncoprotein mouse breast cancer model provides a reliable model for human diseases, Am. J. Pathol., 163 (2003) 2113–2126.

[41] W.J. Muller, E. Sinn, P.K. Pattengale, R. Wallace, P. Leder, Single-step induction of mammary adenocarcinoma in transgenic mice bearing the activated c-neu oncogene, Cell, 54 (1988) 105–115.

[42] A.J. Casbon, D. Reynaud, C. Park, E. Khuc, D.D. Gan, K. Schepers, E. Passegue, Z. Werb, Invasive breast cancer reprograms early myeloid differentiation in the bone marrow to generate immunosuppressive neutrophils, Proc. Natl. Acad. Sci. U. S. A., 112 (2015) E566–575.

[43] S. Picelli, O.R. Faridani, A.K. Bjorklund, G. Winberg, S. Sagasser, R. Sandberg, Full-length RNA-seq from single cells using Smart-seq2, Nat Protoc, 9 (2014) 171–181.

[44] S. Anders, P.T. Pyl, W. Huber, HTSeq--a Python framework to work with high-throughput sequencing data, Bioinformatics, 31 (2015) 166–169.

[45] B. Li, C.N. Dewey, RSEM: accurate transcript quantification from RNA-Seq data with or without a reference genome, BMC Bioinformatics, 12 (2011) 323.

[46] A. Butler, P. Hoffman, P. Smibert, E. Papalexi, R. Satija, Integrating single-cell transcriptomic data across different conditions, technologies, and species, Nat. Biotechnol., 36 (2018) 411–420.

[47] A. Dobin, C.A. Davis, F. Schlesinger, J. Drenkow, C. Zaleski, S. Jha, P. Batut, M. Chaisson, T.R. Gingeras, STAR: ultrafast universal RNA-seq aligner, Bioinformatics, 29 (2013) 15–21.

[48] P. Ewels, M. Magnusson, S. Lundin, M. Kaller, MultiQC: summarize analysis results for multiple tools and samples in a single report, Bioinformatics, 32 (2016) 3047–3048.

[49] Y. Zhou, B. Zhou, L. Pache, M. Chang, A.H. Khodabakhshi, O. Tanaseichuk, C. Benner, S.K. Chanda, Metascape provides a biologist-oriented resource for the analysis of systems-level datasets, Nature communications, 10 (2019) 1523.

[50] J. Reimand, M. Kull, H. Peterson, J. Hansen, J. Vilo, g:Profiler--a web-based toolset for functional profiling of gene lists from large-scale experiments, Nucleic Acids Res, 35 (2007) W193–200.

[51] F.K. Swirski, M. Nahrendorf, M. Etzrodt, M. Wildgruber, V. Cortez-Retamozo, P. Panizzi, J.L. Figueiredo, R.H. Kohler, A. Chudnovskiy, P. Waterman, E. Aikawa, T.R. Mempel, P. Libby, R. Weissleder, M.J. Pittet, Identification of splenic reservoir monocytes and their deployment to inflammatory sites, Science, 325 (2009) 612–616.

[52] F. Leuschner, P.J. Rauch, T. Ueno, R. Gorbatov, B. Marinelli, W.W. Lee, P. Dutta, Y. Wei, C. Robbins, Y. Iwamoto, B. Sena, A. Chudnovskiy, P. Panizzi, E. Keliher, J.M. Higgins, P. Libby, M.A. Moskowitz, M.J. Pittet, F.K. Swirski, R. Weissleder, M. Nahrendorf, Rapid monocyte kinetics in acute myocardial infarction are sustained by extramedullary monocytopoiesis, J. Exp. Med., 209 (2012) 123–137.

[53] C.S. Robbins, A. Chudnovskiy, P.J. Rauch, J.L. Figueiredo, Y. Iwamoto, R. Gorbatov, M. Etzrodt, G.F. Weber, T. Ueno, N. van Rooijen, M.J. Mulligan-Kehoe, P. Libby, M. Nahrendorf, M.J. Pittet, R. Weissleder, F.K. Swirski, Extramedullary hematopoiesis generates Ly-6C(high) monocytes that infiltrate atherosclerotic lesions, Circulation, 125 (2012) 364–374.

[54] V.E. Baracos, L. Martin, M. Korc, D.C. Guttridge, K.C.H. Fearon, Cancer-associated cachexia, Nat Rev Dis Primers, 4 (2018) 17105.

[55] R.N. Kaplan, S. Rafii, D. Lyden, Preparing the “soil”: the premetastatic niche, Cancer Res., 66 (2006) 11089–11093.

[56] S. Hiratsuka, A. Watanabe, Y. Sakurai, S. Akashi-Takamura, S. Ishibashi, K. Miyake, M. Shibuya, S. Akira, H. Aburatani, Y. Maru, The S100A8-serum amyloid A3-TLR4 paracrine cascade establishes a pre-metastatic phase, Nature cell biology, 10 (2008) 1349–1355.

[57] R.N. Kaplan, B. Psaila, D. Lyden, Bone marrow cells in the ‘pre-metastatic niche’: within bone and beyond, Cancer Metastasis Rev., 25 (2006) 521–529.

[58] I. Keklikoglou, C. Cianciaruso, E. Guc, M.L. Squadrito, L.M. Spring, S. Tazzyman, L. Lambein, A. Poissonnier, G.B. Ferraro, C. Baer, A. Cassara, A. Guichard, M.L. Iruela-Arispe, C.E. Lewis, L.M. Coussens, A. Bardia, R.K. Jain, J.W. Pollard, M. De Palma, Chemotherapy elicits pro-metastatic extracellular vesicles in breast cancer models, Nature cell biology, 21 (2019) 190–202.

[59] H. Mo, Y. Shi, X. Han, S. Zhou, X. He, P. Liu, J. Yang, C. Zhang, S. Yang, Y. Qin, L. Gui, Y. Shi, J. Yao, L. Zhao, S. Zhang, Absolute monocyte count is a prognostic indicator in a patient with diffuse large B-cell lymphoma after autologous peripheral blood stem cell transplant, Leuk. Lymphoma, 56 (2015) 515–517.

[60] L. Cassetta, K. Bruderek, J. Skrzeczynska-Moncznik, O. Osiecka, X. Hu, I.M. Rundgren, A. Lin, K. Santegoets, U. Horzum, A. Godinho-Santos, G. Zelinskyy, T. Garcia-Tellez, S. Bjelica, B. Taciak, A.O. Kittang, B. Hoing, S. Lang, M. Dixon, V. Muller, J.S. Utikal, D. Karakoc, K.B. Yilmaz, E. Gorka, L. Bodnar, O.E. Anastasiou, C. Bourgeois, R. Badura, M. Kapinska-Mrowiecka, M. Gotic, M. Ter Laan, E. Kers-Rebel, M. Krol, J.F. Santibanez, M. Muller-Trutwin, U. Dittmer, A.E. de Sousa, G. Esendagli, G. Adema, K. Lore, E. Ersvaer, V. Umansky, J.W. Pollard, J. Cichy, S. Brandau, Differential expansion of circulating human MDSC subsets in patients with cancer, infection and inflammation, J Immunother Cancer, 8 (2020).

[61] T. Satoh, K. Nakagawa, F. Sugihara, R. Kuwahara, M. Ashihara, F. Yamane, Y. Minowa, K. Fukushima, I. Ebina, Y. Yoshioka, A. Kumanogoh, S. Akira, Identification of an atypical monocyte and committed progenitor involved in fibrosis, Nature, 541 (2017) 96–101.

[62] V. Cortez-Retamozo, M. Etzrodt, A. Newton, P.J. Rauch, A. Chudnovskiy, C. Berger, R.J. Ryan, Y. Iwamoto, B. Marinelli, R. Gorbatov, R. Forghani, T.I. Novobrantseva, V. Koteliansky, J.L. Figueiredo, J.W. Chen, D.G. Anderson, M. Nahrendorf, F.K. Swirski, R. Weissleder, M.J. Pittet, Origins of tumor-associated macrophages and neutrophils, Proc. Natl. Acad. Sci. U. S. A., 109 (2012) 2491–2496.

[63] V. Cortez-Retamozo, C. Engblom, M.J. Pittet, Remote control of macrophage production by cancer, Oncoimmunology, 2 (2013) e24183.

[64] H.O. Smith, P.S. Anderson, D.Y. Kuo, G.L. Goldberg, C.L. DeVictoria, C.A. Boocock, J.G. Jones, C.D. Runowicz, E.R. Stanley, J.W. Pollard, The role of colony-stimulating factor 1 and its receptor in the etiopathogenesis of endometrial adenocarcinoma, Clin. Cancer Res., 1 (1995) 313–325.

[65] R. Tang, F. Beuvon, M. Ojeda, V. Mosseri, P. Pouillart, S. Scholl, M-CSF (Monocyte Colony Stimulating Factor) and M-CSF Receptor Expression by Breast Tumor Cells: M-CSF Mediated Recruitment of Tumour Infiltrating Monocytes?, J. Cell. Biochem., 50 (1992) 350–3560.

[66] X. Wu, V.L. Dao Thi, Y. Huang, E. Billerbeck, D. Saha, H.H. Hoffmann, Y. Wang, L.A.V. Silva, S. Sarbanes, T. Sun, L. Andrus, Y. Yu, C. Quirk, M. Li, M.R. MacDonald, W.M. Schneider, X. An, B.R. Rosenberg, C.M. Rice, Intrinsic Immunity Shapes Viral Resistance of Stem Cells, Cell, 172 (2018) 423–438 e425.

