## Supplementary Figures for "Systemic influences of mammary cancer on monocytes in mice"

**Supplementary Figure Legends**

**Supplementary Figure 1**: Flow cytometric gating and cell-sorting strategy for blood monocytes and neutrophils used in Figure 1, 2, 4 and 6.

**Supplementary Figure 2**: Flow cytometric gating strategy for BM monocytes in Figure 1 and Figure 2E.

**Supplementary Figure 3**: Flow cytometric gating strategy for identification and sorting of BM Ly6C$^{high}$ monocytes, MDPs and cMoPs in Figure 2A, C and Figure 6.

**Supplementary Figure 4**: Flow cytometric gating strategy for identification and sorting of BM LK in Figure 2A and B.

**Supplementary Figure 5:** Flow cytometric gating strategy for identification of BM myeloid progenitors in Figure 2D.

**Supplementary Figure 6:** Flow cytometric gating strategy for identification of splenic Ly6C$^{high}$ monocytes and MDP in Figure 3.

**Supplementary Figure 7**: Identification of BrdU$^{+}$ monocytes in the blood (A), BM (B) and spleen (C) in Figure 2D-F and Figure 3F. Example staining taken at 24 (A) or 1 (B-C) hrs post-injection of BrdU.

**Supplementary Figure 8:** Bulk RNA sequencing of Ly6C$^{high}$ and Ly6C$^{low}$ blood monocytes from C57BL/6 mice with late cancer and littermate controls. *(A) PCA of all monocytic samples derived from total RNAseq (B) PCA of Ly6C$^{low}$ monocytes derived from total RNAseq*

### Supplementary Figure 1

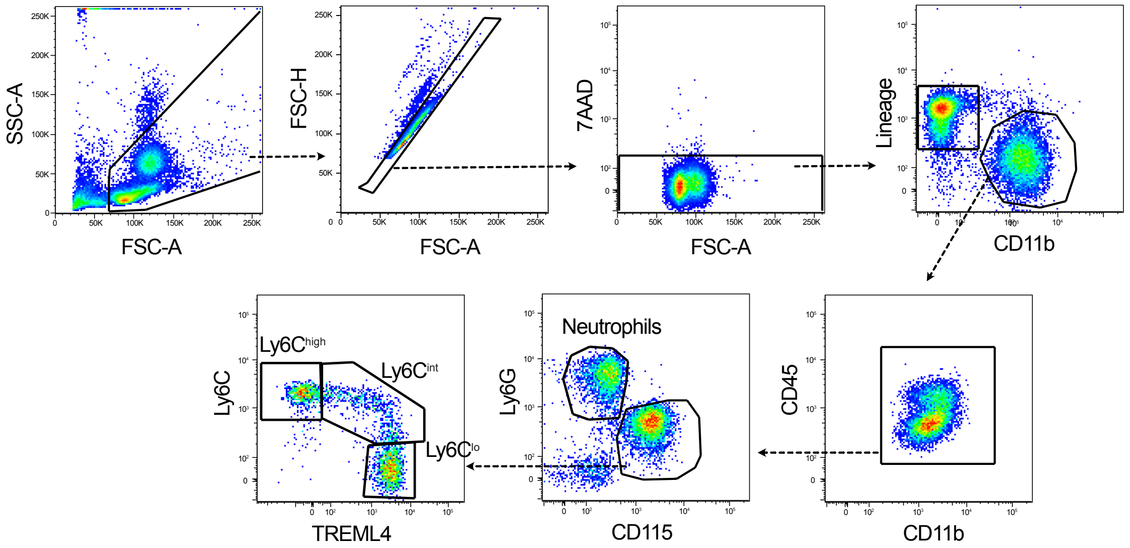

**Supplementary Figure 2**

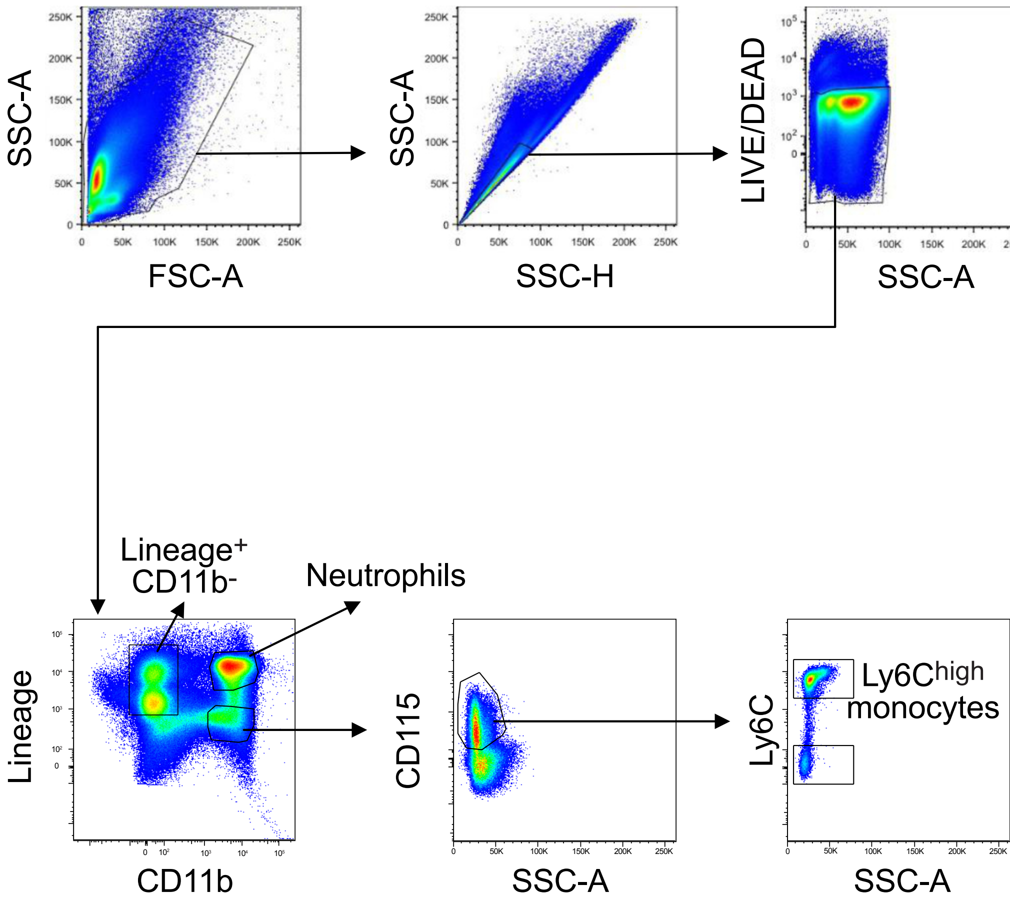

### Supplementary Figure 3

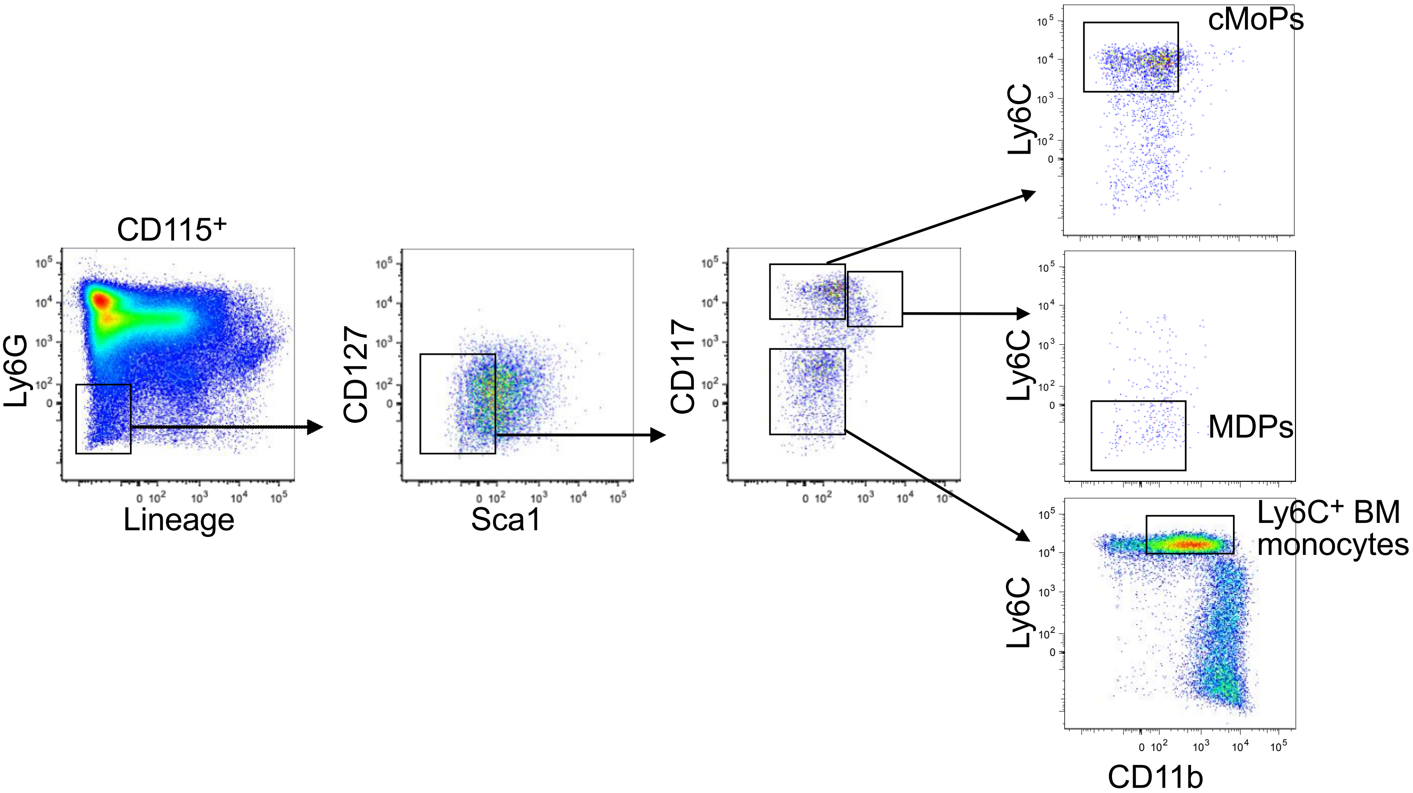

**Supplementary Figure 4**

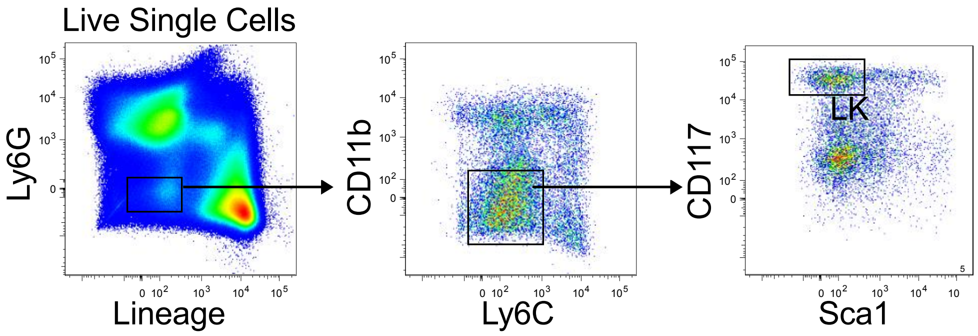

**Supplementary Figure 5**

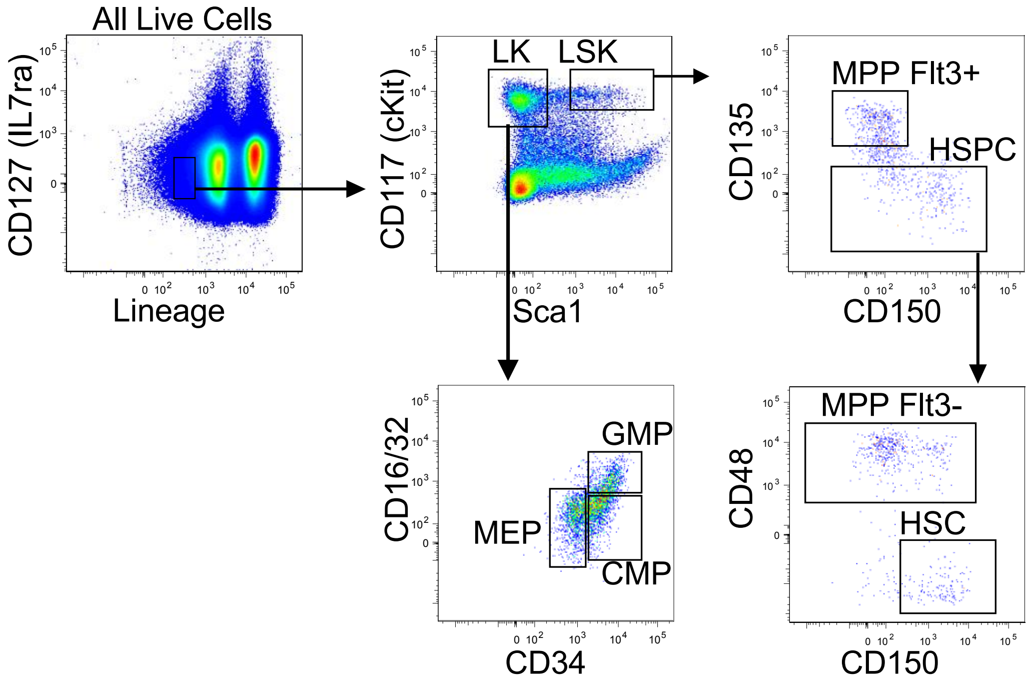

**Supplementary Figure 6**

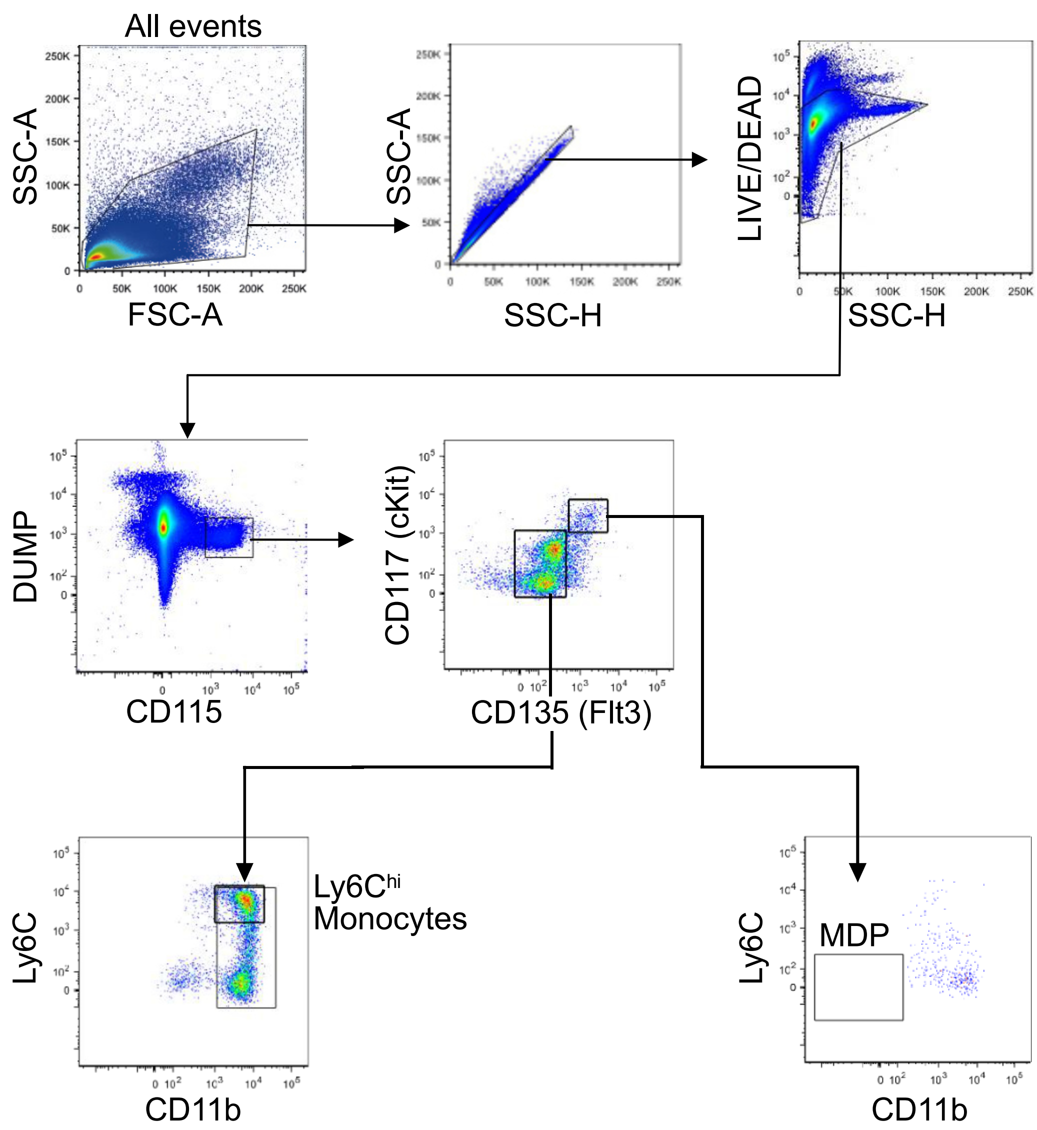

**Supplementary Figure 7**

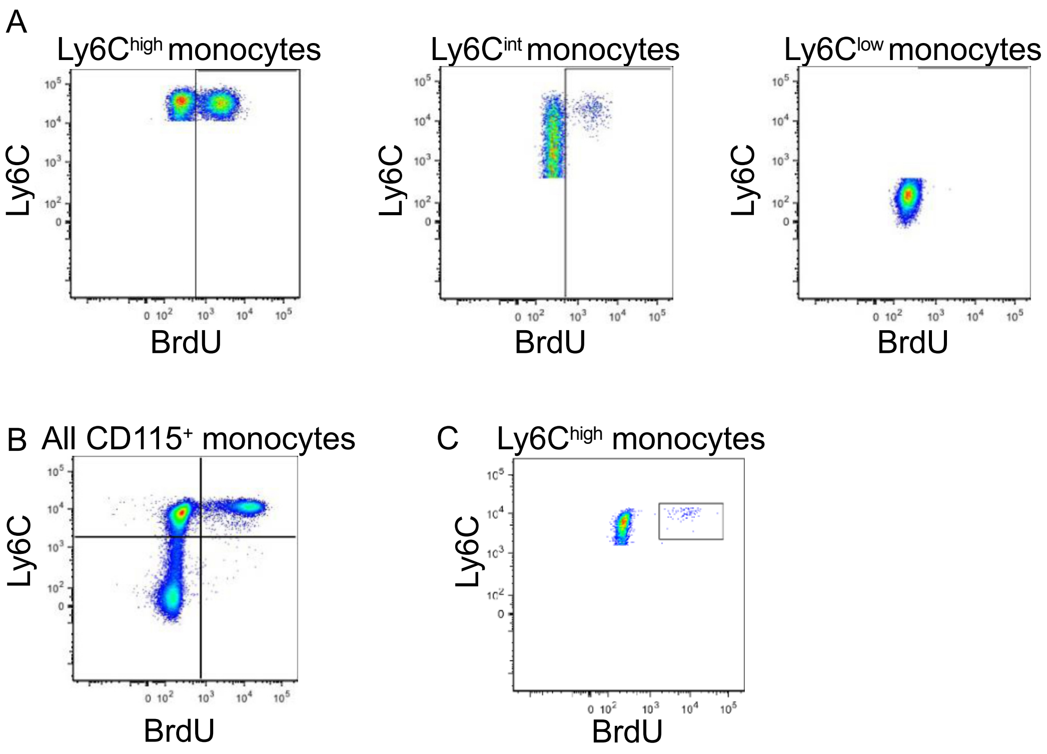


`

### Supplementary Figure 8

**A**

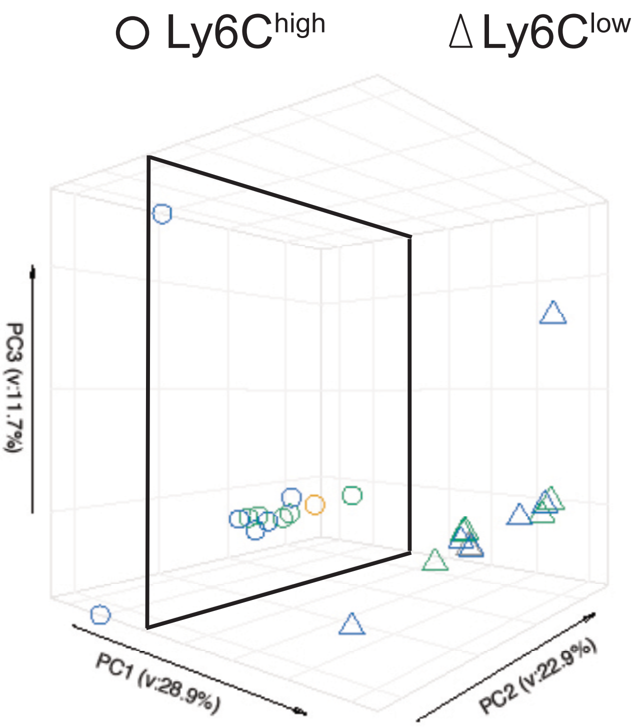


**B**

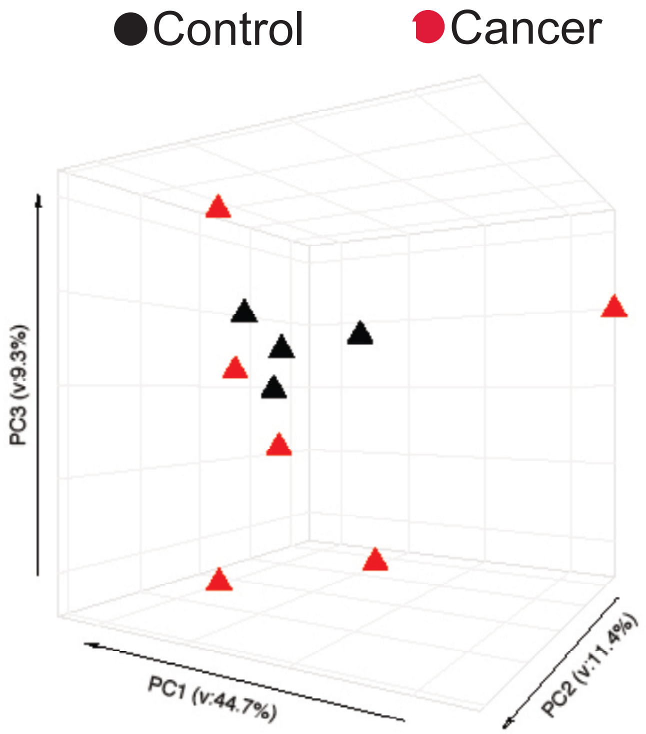
